# Effects of intergenerational transmission of small intestinal bacteria cultured from stunted Bangladeshi children with enteropathy

**DOI:** 10.1101/2024.11.01.621574

**Authors:** Kali M. Pruss, Clara Kao, Alexandra E. Byrne, Robert Y. Chen, Blanda Di Luccia, Laura Karvelyte, Reyan Coskun, Mackenzie Lemieux, Keshav Nepal, Daniel M. Webber, Matthew C. Hibberd, Yi Wang, Haoxin Liu, Dmitry A. Rodionov, Andrei L. Osterman, Marco Colonna, Christian Maueroder, Kodi Ravichandran, Michael J. Barratt, Tahmeed Ahmed, Jeffrey I. Gordon

## Abstract

Environmental enteric dysfunction (EED), a small intestinal disorder found at a high prevalence in stunted children, is associated with gut mucosal barrier disruption and decreased absorptive capacity^1–4^. To test the hypothesis that intergenerational transmission of a perturbed small intestinal microbiota contributes to undernutrition by inducing EED^5^, we characterized two consortia of bacterial strains cultured from duodenal aspirates from stunted Bangladeshi children with EED – one of which induced local and systemic inflammation in female gnotobiotic mice. Offspring of dams colonized with the inflammatory consortium exhibited impaired prenatal and postnatal growth, as well as immunologic changes phenocopying features of EED in children. Dam-to-pup transmission of the inflammatory consortium produced, in recently weaned offspring, alterations in inter-cellular signaling pathways related to intestinal epithelial cell renewal, barrier integrity and immune function. Cohousing of mice harboring the inflammatory or non-inflammatory consortia and subsequent screening of candidate disease-promoting bacterial isolates identified *Campylobacter concisus,* an organism typically found in the oral microbiota, as a contributor to enteropathy. The *C. concisus* strain induced, in a host nitric oxide synthase (NOS)-dependent manner, pro-inflammatory cytokine signaling. Moreover, host-derived nutrients generated by NOS augmented *C. concisus* growth. This preclinical model should facilitate identification of small intestinal microbiota-targeted therapeutics for (intergenerational) undernutrition.

---

Undernutrition is a significant global health challenge. Numerous epidemiologic studies indicate that stunting in mothers is associated with low birth weight and postnatal linear growth faltering (stunting) in their offspring^6,7^. Stunting in infants and children is accompanied by increased risk of infection, metabolic/hormonal imbalances later in life^8^, as well as neurodevelopmental/cognitive abnormalities^9–11^.

One hypothesis is that a perturbed small intestinal microbiota^2^ contributes to undernutrition by inducing environmental enteric dysfunction (EED)^1,12–14^ and subsequent transmission of the maternal small intestinal microbiota to offspring perpetuates intergenerational EED^5^. Evidence that the small intestinal microbiota contributes to the pathogenesis of EED comes in part from the Bangladeshi EED (BEED) study, in which stunted children who had failed to respond to a nutritional intervention underwent esophago-gastroduodenoscopy (EGD)^2,15^. Almost all (95%) of these children displayed histopathologic evidence of EED^2^. Sequencing bacterial 16S rRNA amplicons generated from duodenal aspirates collected during EGD revealed a ‘core’ group of 14 taxa (amplicon sequence variants, ASVs) that were present in >80% of the children. The absolute abundances of these taxa, not typically classified as enteropathogens, were significantly negatively correlated with linear growth (length-for-age Z score, LAZ) and positively correlated with levels of duodenal mucosal proteins involved in immunoinflammatory responses^2^.

Deciphering the role of the small intestinal microbiota in the pathogenesis of EED has been hampered by several factors. For example, EGD is limited to the proximal intestine and carries potential risks, precluding, for ethical reasons, its use in healthy children to define what constitutes a ‘normal’ small intestinal microbiota. In addition, the volume of fluid and biomass of microbes that can be retrieved from the lumen of the small intestine with EGD is small, limiting the number of ways the material can be employed for downstream analyses. Moreover, preclinical studies are necessary to define mechanisms and establish causal relationships between members of the microbiota and EED. Current mouse models rely on either provision of a low-protein diet, which has not been explicitly defined as a causative factor for EED in humans, and/or administration of a single pathogen or a broad inflammatory insult (e.g. treatment with LPS)^16–19^. While these models are valuable for understanding effects of protein restriction on immunity and the virulence mechanisms of enteric pathogens, they do not simulate the complexity of small intestinal bacterial community dynamics and interactions with the host that likely play a role in pathophysiology of EED in humans.

In the current study, we sought to model maternal-to-offspring transmission of different small intestinal bacterial consortia from children in the BEED study that elicit discordant intestinal and systemic inflammatory responses in gnotobiotic mice. Our goal was to use these consortia to obtain insights about host cellular and molecular responses and to identify bacteria that have a causal role in pathology associated with EED.

## Results

### Development of inflammatory and non-inflammatory EED-derived bacterial consortia

In a previous pilot study^2^, a collection of 184 bacterial isolates cultured from the duodenal aspirates of children in the BEED study were pooled and introduced by oral gavage directly into adult germ-free mice fed a diet formulated based on foods consumed by children living in Mirpur, Bangladesh, the urban slum where the clinical study was performed (‘Mirpur-18’ diet, **Supplementary Table 1a**). Compared to controls gavaged with intact cecal contents from conventionally-raised mice (conventionalized animals, CONV-D), the duodenal aspirate-derived culture collection induced an enteropathy characterized by (i) patchy immunoinflammatory infiltrates in the small intestine, (ii) increased duodenal crypt depth, (iii) reduced expression of tight junction proteins and increased expression of genes involved in anti-microbial defense (*Reg3β* and *Reg3γ*) in the duodenum, and (iv) elevated levels of matrix metalloproteinase 8 (MMP8) in serum as well as along the length of the small intestine^2^.

We sought to reduce the complexity of the 184-member bacterial consortium derived from the EED donors. To do so, we chose a representative of each bacterial species (based on sequencing full-length 16S rRNA amplicons, **Supplementary Table 1b**) and pooled these 39 isolates into a ‘species-representative subset consortium’. These two consortia were gavaged into separate groups of just-weaned 4-5-week-old germ-free mice (n=5 animals/group) fed the Mirpur-18 diet. Mice colonized with all 184 isolates gained significantly less weight over 28 days than mice colonized with the species-representative subset (**Extended Data Fig. 1a, Supplementary Table 2a**). Serum levels of insulin-like growth factor 1 (IGF-1) were significantly correlated with body mass at 9 and 28 days after colonization across all animals (**Extended Data Fig. 1b**), although there were no statistically significant differences between treatment groups (**Supplementary Table 2b**). Leptin levels were diminished compared to CONV-D animals, but not significantly different between mice that received the 184-member and species-representative consortia (**Extended Data Fig. 1c, Supplementary Table 2b**). Together, these results suggest that additional factors contributed to the reduced weight gain observed in animals that received the complete consortium.

Sequencing of RNA isolated from the duodenums, jejunums and ileums of these animals followed by gene set enrichment analysis (GSEA) revealed that a large proportion of the gene ontology (GO) categories that were significantly enriched (*q*<0.05) between animals colonized with the full versus subset bacterial consortia were related to immune function, including genes involved in the anti-bacterial defense response and leukocyte activation (**Extended Data Fig. 1d,e**). Levels of the protein lipocalin-2 (LCN2/NGAL; neutrophil gelatinase-associated lipocalin) were also significantly elevated in the serum of mice colonized with the full 184-isolate consortium (**Extended Data Fig. 1f**); previously we had observed that LCN2 levels in the duodenal mucosa of children with EED were strongly correlated with the absolute abundances of the 14 duodenal EED-associated ‘core taxa’^2^. Based on the higher levels of intestinal and systemic inflammation in animals colonized with the full consortium of EED-derived bacteria, we named this community ‘child small intestinal inflammation-inducing’ (cSI-I); the species-representative subset was designated ‘child small intestinal non-inflammatory’ (cSI-N).

### Modeling maternal-offspring transmission of EED donor small intestinal bacterial consortia

As a prelude to determining whether inflammation induced by the cSI-I consortium could be transmitted from dams to their offspring, we performed a diet oscillation experiment. Adult germ-free animals fed a standard mouse diet were colonized with either the cSI-I or cSI-N consortium. Mice were maintained on the standard chow for one week, then given a diet designed to be representative of that consumed by adults residing in the Mirpur district of Dhaka (‘Adult Mirpur’ diet; **Supplementary Table 1a**), followed by a return to standard diet for one week. LCN2 levels were measured in serum samples obtained at the end of each of the 7-day diet periods (**Extended Data Fig. 1g**). Statistically significant increases in systemic inflammation were documented in both groups of mice when fed Adult Mirpur compared to standard mouse diet. Moreover, LCN2 levels were significantly higher in mice colonized with cSI-I compared to cSI-N in both diet contexts (**Extended Data Fig. 1h, Supplementary Table 2b**).

Based on these results, germ-free adult female C57Bl/6J mice were placed on the ‘Adult Mirpur’ diet. Three days later, separate groups of animals (n=4 dams/group) received an oral gavage of the cSI-I or cSI-N consortia, or intact cecal contents from conventionally raised adult C57Bl/6J mice (**Fig. 1a, Extended Data Fig. 2a**). One week after colonization, female mice were mated with germ-free males. Pups born to these dams were weaned onto the ‘Mirpur-18’ diet (**Supplementary Table 1a**) which they consumed *ad libitum* until euthanasia on postnatal day 37 (P37). To assess the reproducibility and timing of onset of EED features, second and third litters produced by the same dams were euthanized either on P37 or P14, respectively (**Extended Data Fig. 2a,b**).

**Fig. 1.**
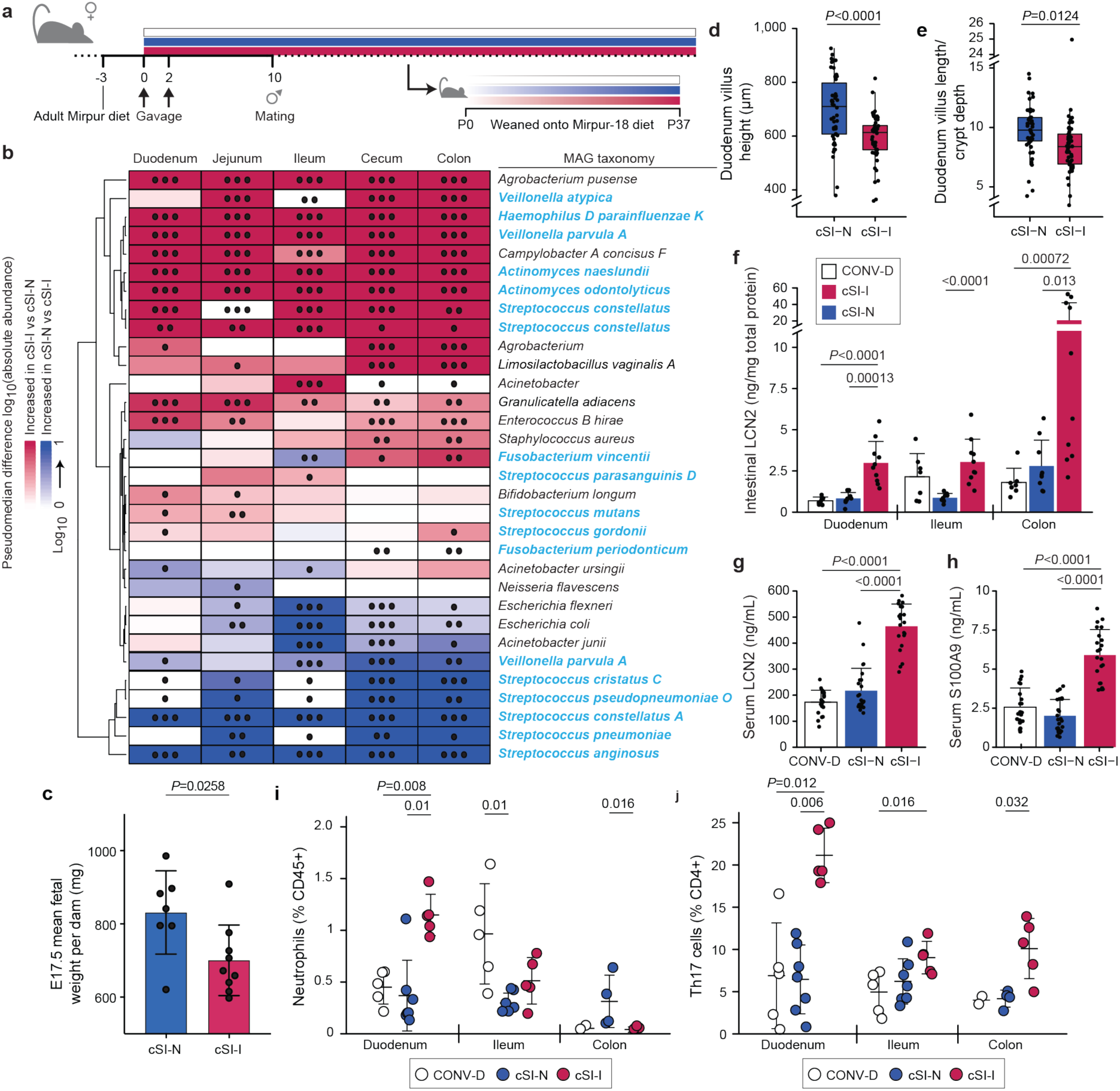
Dam-to-offspring transmission of bacteria cultured from duodenal aspirates collected from Bangladeshi children with EED. **a,** Design of intergenerational transmission experiment. **b,** Differential abundances of bacterial MAGs (rows) along the length of the intestine (columns) of P37 pups born to dams harboring cSI-N or cSI-I bacterial consortia (n=21-25 mice/group). MAGs were included in the heatmap if they demonstrated significant differential log_10_ absolute abundance in at least one of the intestinal locations. MAG taxonomy colored in blue indicates correspondence to ASVs representing ‘core taxa’ identified in the small intestines of stunted children with EED in the BEED study. •, *P*-adj < 0.05; ••, *P*-adj < 0.01; •••, *P*-adj < 0.001, FDR-corrected Wilcoxon rank-sum tests. **c,** Average fetal mass for a given dam at E17.5. Dams were colonized with the cSI-I or cSI-N consortia for two weeks prior to mating (n=7-9 dams/group; n=2-10 fetuses/litter, unpaired t-test). **d,e,** Villus length (**d**), and ratio of villus length to crypt depth (**e**) in the duodenum of P37 cSI-I compared to cSI-N mice (n=5 animals/group, boxes denote interquartile range). **f,** Levels of lipocalin-2 (LCN2) protein in intestinal tissue from P37 mice (n=7-10 mice/group). **g,h,** Serum levels of LCN2 (**g**), and S100A9 protein (**h**), in P37 pups (n=21-25 mice/group). For c-e, n=4-6 litters/group, 3-7 pups/litter. **i,j,** Frequency of neutrophils (**i**), and Th17 cells (**j**), along the length of the gut (n=5-7 mice/group, each point represents an individual animal). For **d-j,** statistics shown are the result of Wilcoxon-rank sum tests. For **c-j**, mean values ± s.d. are shown.

We identified bacterial members of each consortium that successfully colonized recipient dams and were transmitted to their offspring using long-read shotgun DNA sequencing of cecal contents from dams and P37 pups followed by assembly of metagenome-assembled genomes (MAGs) (n=4 dams and n=22-25 offspring/consortium). We reasoned that this approach would be useful for initial characterization of colonizing members of culture collections where multiple strains of a given species may exist. A total of 51 unique high-quality MAGs (defined as ≥85% complete and ≤5% contaminated based on marker gene analysis, **Supplementary Table 1c**) were generated from cecal contents of dams and P37 offspring harboring either of the two cSI consortia. A total of 25 MAGs from the cSI-I consortium colonized dams and all were represented in their P37 offspring (**Extended Data Fig. 2c**). Twenty-eight MAGs from the cSI-N consortium colonized dams; all 28 also colonized their P37 offspring (threshold criteria for colonization: >0.0001% relative abundance in at least one intestinal segment in at least 50% of animals, **Supplementary Table 1d, Extended Data Fig. 2c**). Based on short-read shotgun sequencing-based calculations of MAG abundances in fecal samples collected from dams, the cSI-I and cSI-N communities remained stable between 4 and 17 weeks after gavage of these cultured consortia even if the animals had one or more litters during this period (*P*=0.09, PERMANOVA; n=4 dams/group; **Extended Data Fig. 2d,e**).

The absolute abundances of 21 MAGs were significantly higher in at least one intestinal segment of P37 offspring colonized with the cSI-I consortium compared to cSI-N; the taxonomy of 12 of these 21 MAGs corresponded to ASVs we previously identified as ‘core taxa’ present in >80% of the duodenal microbiota of Bangladeshi children with duodenal mucosal biopsy-diagnosed EED (**Fig. 1b**)^2^. Moreover, analysis of fecal samples collected from the diet oscillation experiment described in **Extended Data Fig. 1g**, disclosed that 19 of these 21 MAGs were significantly higher in cSI-I compared to cSI-N adult mice when fed either Adult Mirpur or standard mouse diets (**Extended Data Fig. 2f,g**). Fewer MAGs were detected in P14 offspring (88% of MAGs that colonized cSI-I dams; 68% of cSI-N MAGs that colonized dams). Of the 21 MAGs with significantly higher absolute abundance in P37 cSI-I offspring compared to their cSI-N counterparts, seven were significantly higher in the cecum and/or colon of P14 offspring (**Extended Data Fig. 3a**).

### Reduced prenatal and postnatal growth with cSI-I consortium

We colonized a separate group of adult female mice with the cSI-I or cSI-N consortia two weeks before mating. Pregnant dams were euthanized when placentation just completed (embryonic day 11.5, E11.5) or near the end of gestation (E17.5). Fetal weights were significantly lower in cSI-I compared to cSI-N dams at E17.5 (**Fig. 1c,** n=7-9 dams/group; n=2-10 fetuses/litter), but not at E11.5 (n=9-10 dams/group; n=4-8 fetuses/litter) (**Extended Data Fig. 3b**; **Supplementary Table 2a**). Placental weights were not significantly different between treatment groups at either timepoint (**Extended Data Fig. 3c,d, Supplementary Table 2a**).

Body mass (**Extended Data Fig. 3e**) and serum levels of IGF-1 (**Extended Data Fig. 3f**) were significantly reduced in pre-weaning (P14) offspring of cSI-I dams compared to their cSI-N counterparts (**Supplementary Table 2a,b**). P14 cSI-I pups displayed significantly higher serum levels of the pro-inflammatory proteins LCN2, S100A9 (S100 calcium binding protein A9), CHI3L1 (Chitinase-3-like protein 1), MMP8 (matrix metallopeptidase 8, or neutrophil collagenase), as well as the immune cell chemoattractant CXCL1 (chemokine ligand 1) compared to their cSI-N and CONV-D counterparts (**Extended Data Fig. 3g-j**; **Supplementary Table 2b**). LCN2, CHI3L1, S100A9 and MMP8 belong to a group of inflammatory proteins whose expression in duodenal mucosal biopsies was significantly positively correlated with the abundances of the ‘core’ EED-associated bacterial taxa in Bangladeshi children with EED^2^. LCN2, S100A9 and CHI3L1 were similarly elevated in duodenal, ileal and colonic tissue of cSI-I animals (**Extended Data Fig. 3k-m, Supplementary Table 2b**). Thus, systemic and intestinal inflammation were already evident in these animals prior to completion of weaning.

### cSI-I dams and their offspring exhibit enteropathy

Postnatal day 14 represents a critical period of maturation of lymphoid organs and the adaptive immune system. To determine whether the effects of the cSI-I and cSI-N consortia persisted through the weaning period, we assessed offspring on P37, which represents a developmental state more akin to adolescents^20^. Children with EED have villus blunting: histomorphometric analyses of the small intestines of P37 offspring of cSI-I dams revealed significantly diminished duodenal villus height (**Fig. 1d**) and a significantly diminished ratio of villus height to crypt depth in the duodenum and ileum compared to their cSI-N counterparts (**Fig. 1e, Extended Data Fig. 4a**; **Supplementary Table 2c**). Crypt depth in the ileum of P37 cSI-I mice trended towards increased relative to cSI-N animals (*P*=0.09, Wilcoxon rank-sum). The observed increase in crypt depth is consistent with the notion that there is a proliferative response involving stem and transit-amplifying cells in this model, but one that is insufficient to restore villus height (see ‘***Proliferative signaling to stem and transit-amplifying cells’*** below).

LCN2 levels in duodenal, ileal and colonic tissue, as well as serum, were significantly elevated in P37 mice born to dams harboring the cSI-I compared to the cSI-N consortium, or to conventionalized (CONV-D) dams (**Figs. 1f,g**). S100A9, MMP8, CXCL1 and IL-17 were similarly elevated in the serum of P37 cSI-I offspring (**Fig. 1h**, **Extended Data Fig. 4b,c**; **Supplementary Table 2b**). As in P14 animals, duodenal, ileal and colonic tissue levels of S100A9 and CHI3L1 were significantly higher in P37 offspring of cSI-I dams (**Extended Data Fig. 4d,e**; **Supplementary Table 2b**). There were no statistically significant differences in the serum levels of these proteins between litters, nor between male and female offspring in the same treatment group (*P*>0.05, Wilcoxon rank-sum tests).

Compared to P37 cSI-N and CONV-D mice, serum levels of Dickkopf-1 (DKK1), Osteoprotegerin (OPG) and Fibroblast Growth Factor-23 (FGF23) were significantly higher in cSI-I animals, indicating increased osteocyte and bone remodeling activity (*P*<0.015 for all comparisons, Wilcoxon rank-sum test; **Supplementary Table 2b**). Micro-computed tomography of femurs disclosed significantly increased cortical tissue mineral density in P37 cSI-I versus cSI-N animals (*P*=0.03, Wilcoxon rank-sum test; n=19-21 mice/group); no significant differences in other cortical and trabecular parameters were found between the three treatment groups (**Supplementary Table 2d**).

#### Small intestinal tissue acylcarnitines

Plasma acylcarnitines are elevated in a fasting state and have been identified as a biomarker of EED in children^21^. Moreover, fecal acylcarnitines are elevated in the dysbiotic microbial states associated with inflammatory bowel disease^22^ and can activate pro-inflammatory signaling cascades^23,24^. We quantified acylcarnitine levels in intestinal tissue instead of plasma to directly examine whether differences in host fatty acid metabolism were produced as a result of colonization with the cSI-I compared to the cSI-N consortium. Several medium and long-chain acylcarnitines were significantly elevated in the intestine of cSI-I compared to cSI-N or CONV-D offspring (**Extended Data Fig. 4f, Supplementary Table 2e**), suggesting that the carnitine shuttle may be impaired in the intestine.

#### Flow cytometry of immune cell populations in the intestinal lamina propria

The frequency of neutrophils was significantly increased in the duodenum of P37 cSI-I offspring, but not in the ileum or colon (**Fig. 1i**). CD3^+^ and CD4^+^ cells were also higher in the duodenum, ileum and colon of cSI-I animals compared to age-matched cSI-N or CONV-D mice (**Extended Data Fig. 4g,h**). Among CD4^+^ cells, the Th17 population was significantly higher in the duodenum, ileum and colon of cSI-I offspring (**Fig. 1j**), whereas Th1 cells were diminished in their duodenum and Tregs were diminished in their duodenum and colon (**Supplementary Table 2f**); these findings lead us to speculate that a pro-inflammatory cytokine milieu in the intestines of P37 cSI-I offspring promotes Th17 cell differentiation and proliferation.

These protein biomarker, neutrophil, and T cell phenotypes were also evident in dams: the frequencies of duodenal neutrophils, CD3^+^ cells, CD4^+^ T cells and Th17 cells were all significantly elevated in cSI-I adult female mice. Th1 cells and Tregs were similarly significantly reduced in cSI-I compared to cSI-N mothers (*P*<0.05 for all comparisons, Wilcoxon rank-sum test; n=5 mice/group; **Supplementary Table 2f**). The observed increases in CD3^+^ cells and CD4^+^ T cells in the intestinal tissues of dams and their offspring harboring cSI-I compared to cSI-N consortia are consistent with prior characterization of EED in children as a T-cell mediated enteropathy^13,25^.

### Intestinal epithelial cellular responses to cSI-I and cSI-N consortia

#### Single nucleus RNA-sequencing (snRNA-Seq)

To characterize the responses of different intestinal epithelial cell types to the EED donor-derived bacterial consortia, we performed snRNA-Seq on intact segments of frozen duodenum and ileum harvested from P37 cSI-I and cSI-N mice. A total of 15,135 duodenal nuclei (**Fig. 2a**) and 20,584 ileal nuclei (**Fig. 2b**) collected from 3 littermates/group passed our quality metrics (see *Methods*). Analysis of marker gene expression identified nuclei from epithelial lineages (stem and transit amplifying cells and their enterocyte, goblet, Paneth, enteroendocrine and tuft cell descendants), as well as from mesenchymal lineages (endothelial, smooth muscle, neuronal, immune and interstitial cells of Cajal). Enterocytes comprised the largest proportion of nuclei (**Supplementary Table 3b**) and based on marker gene expression were divided into villus base, mid-villus and villus tip subpopulations. A significantly higher proportion of Paneth cells were present in the duodenum of cSI-I offspring than their cSI-N counterparts (**Supplementary Table 3b**). Increased density^26^ of Paneth cells as well as degranulation^27^ have both been reported in Zambian children with EED.

‘Pseudo-bulk’ analysis of differential gene expression in enterocytes positioned at different locations along the crypt-villus axis in both the duodenum and ileum revealed an enhanced epithelial immune response in P37 cSI-I compared to cSI-N mice. Of the 133 GO categories significantly enriched in at least two subpopulations of enterocytes in cSI-I offspring, 35 were involved in bacterial recognition and defense, as well as recruitment/migration and activation of immune cells (**Extended Data Fig. 5a, Supplementary Table 3c**).Epithelial homeostasis requires a balance of differentiation, programmed cell death, and clearance of dying cells and cellular debris. This process, termed ‘efferocytosis’, involves recognition and clearance of dying cells by phagocytes, including both professional phagocytes and healthy neighboring intestinal epithelial cells^28–30^. Efferocytosis can be viewed as a process that sits at the interface between regulation of gut epithelial renewal, inflammation and the microbiota^31^. Based on published literature^32–34^, we manually curated an ‘efferocytosis’ gene set composed of 273 genes. This gene set was significantly enriched in cSI-I mid-villus duodenal enterocytes (*P*=0.019, **Extended Data Fig. 5b**). Genes enriched in cSI-I duodenal enterocytes included caspases linked to apoptosis (*Casp3*), genes linked to phosphatidylserine exposure during apoptosis (*Xkr8, ATP11c*), a channel that regulates apoptotic cell communication with the microbiota (*Panx1*), several efferocytotic receptors/recognition mediators (*Mertk, Stab1*, *Mfge8*), as well as cytosolic proteins linked to intracellular signaling during efferocytosis (*Elmo1, Elmo2, Dock1, Crk*, see **Supplementary Table 3c**). These findings suggest that apoptosis and subsequent efferocytosis within the small intestine play a role in shaping the epithelial response to the cSI-I consortium.

**Fig. 2.**
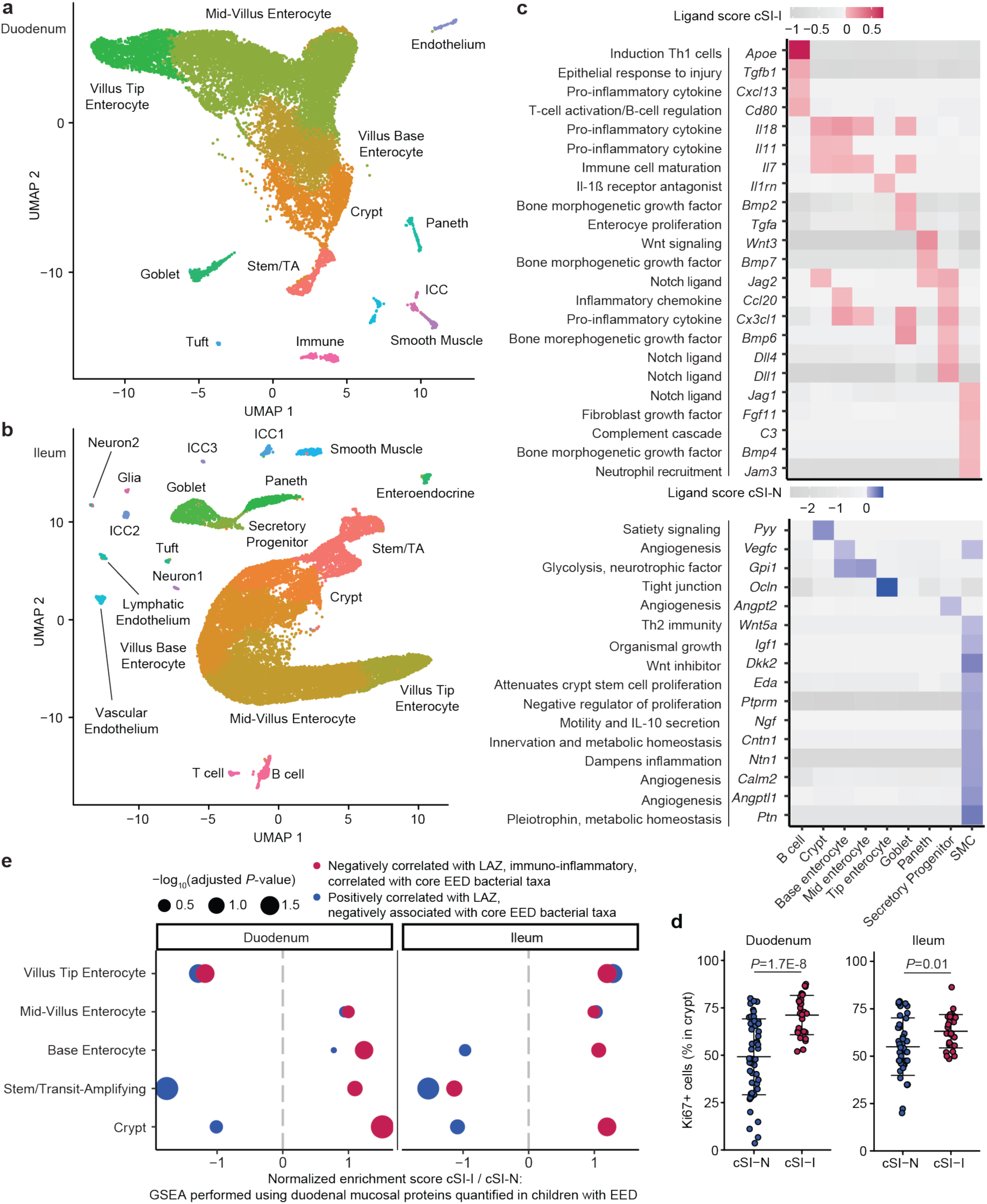
Increased proliferative signaling in the small intestine of P37 offspring of cSI-I dams. **a,b,** Uniform manifold and projection (UMAP) plot of single nuclei isolated from the duodenum (**a**), and ileum (**b**) of P37 offspring (n=3 mice/group). **c,** Intercellular signaling from cell populations (columns) to stem/transit amplifying (TA) cells in the ileum. A subset of ligands identified by *NicheNet* are shown that were significantly differentially expressed between cSI-I and cSI-N animals. Annotations for these ligands (left) are based on literature findings. Their paired receptors are shown in **Extended Data Fig. 6a**. Ligands that were more highly expressed in cSI-I mice are shown in red (top of panel); those more highly expressed in cSI-N animals are shown in blue (bottom of panel). **d,** Percent of all cells in a crypt that were Ki67^+^ in the duodenum (left) or ileum (right) (n=3-5 mice/group; each dot represents a single crypt-villus unit; 10 crypts analyzed per intestinal segment per mouse). *P*-values were determined by Tukey’s post-hoc tests; mean values ± s.d. are shown. **e,** Gene set enrichment analysis (GSEA) along the crypt-villus axis, focusing on genes encoding duodenal mucosal proteins whose levels were quantified in children with EED in the BEED study.

#### Proliferative signaling to stem and transit-amplifying cells

To assess the effects of the SI consortia on epithelial regeneration, we used the algorithm *NicheNet*^35,36^ to characterize intercellular signaling in cSI-I versus cSI-N P37 offspring. *NicheNet* surveys expression of known protein ligands by user-designated ‘sender’ cell types and their cognate receptors within a ‘receiver’ population. For our analysis, we defined stem and transit amplifying (stem/TA) cells as ‘receivers’ and all other cell types within the SI as ‘senders’ (**Fig. 2c**). Bone morphogenetic protein (Bmp), Wnt and Notch signaling are three of the major pathways that mediate cellular proliferation during normal intestinal development, responses to injury and hyperplastic inflammatory responses^37,38^. In the ileum, ligands more highly expressed in cSI-I compared to cSI-N mice included *Wnt3* in Paneth cells (and their receptors *Lrp6, Ryk* and *Bmpr1a* in TA/stem cells, **Extended Data Fig. 6a, Supplementary Table 3c**), as well as Paneth cell *Bmp7* (and its *Bmpr1a* and *Bmpr2* receptors in stem/TA cells). *Bmp6* and *Bmp2,* two other ligands for *Bmpr1a* and *Bmpr2* receptors, were also expressed at higher levels in ileal cSI-I goblet cells (**Fig. 2c, Extended Data Fig. 6a**). Expression of *Wnt5a* and the Wnt inhibitor *Dkk2* were significantly diminished in ileal smooth muscle cells (SMCs) in cSI-I mice, as were their receptors in stem/TA cells (*Cftr* and *Ptprk* in the case of *Wnt5a* and *Lrp6* in the case of *Dkk2*) (**Fig. 2c, Extended Data Fig. 6a, Supplementary Table 3c**). SMC-derived *Wnt5a* augments Th2 immunity and inhibits cellular proliferation by downregulating expression of *Ctnnb1* (ß-catenin)^39^. Quantification of Ki67, a marker of cellular proliferation, in duodenal and ileal crypts provided additional evidence of increased epithelial regeneration in cSI-I animals (**Fig. 2d, Extended Data Fig. 6b**; **Supplementary Table 2c**).

#### Contextualizing intestinal transcriptional responses with proteomic features in duodenal mucosa of children with EED

To further assess the degree to which the transcriptional response in enterocytes present in the duodenum and ileum of P37 offspring recapitulates pathophysiologic changes we had documented in the upper gastrointestinal tract of Bangladeshi children with EED^2^, we performed GSEA with mouse homologs of duodenal mucosal proteins that were: (i) negatively correlated with length-for-age-Z-score (LAZ) and positively correlated with the levels of core EED bacterial taxa in BEED study participants, or (ii) positively correlated with LAZ in these children and negatively associated with core EED taxa (**Fig. 2e**). Expression of mouse genes homologous to the human duodenal proteins that were negatively associated with growth and positively correlated with the absolute abundances of core taxa were enriched in cSI-I enterocytes in the duodenum (crypt through mid-villus) and in the ileum (crypt and along the length of the villus) (**Fig. 2e**). Genes encoding antimicrobial peptides (*Reg3β* and *Reg3γ*), the pro-inflammatory cytokine *Il18*, the heat-shock protein *Hsp90aa1* and the immune cell chemoattractant *Ccl28* were more highly expressed in all duodenal enterocyte cell types whereas the type-2 immune response regulator *Arg-2* was more highly expressed in cSI-N offspring (**Extended Data Fig. 6c, Supplementary Table 3c**). Together, these results provide evidence that SI enterocytes play an important role in mediating the immune response in offspring of dams colonized with the cSI-I consortium.

### Cohousing to nominate pathology-inducing bacterial strains

To identify bacterial taxa in the two consortia that either confer or ameliorate the immunoinflammatory state described above, we gavaged separate groups of 4-5-week-old germ-free C57Bl/6J mice consuming the Mirpur-18 diet with the cSI-I or cSI-N culture collections. After nine days, half of the animals from each group were cohoused. This experimental group and the non-cohoused cSI-I and cSI-N control groups were then followed for an additional nine days (n=6 or 8 mice/group/experiment, 2 independent experiments; **Fig. 3a**). Cohoused mice and non-cohoused cSI-I controls gained significantly less weight compared to non-cohoused cSI-N controls (linear mixed effects model; **Supplementary Table 2b**). Irrespective of the initial colonization state, serum concentrations of LCN2 and CHI3L1 were significantly higher in cohoused animals compared to non-cohoused cSI-N controls and did not differ significantly from levels of these biomarkers in non-cohoused cSI-I animals (**Fig. 3b, Supplementary Table 2b**). Levels of LCN2 and CHI3L1 were also significantly higher in colonic tissue and trended higher in duodenal tissue of cohoused mice compared to non-cohoused cSI-N controls (**Fig. 3c**; **Supplementary Table 2b**).

**Fig. 3.**
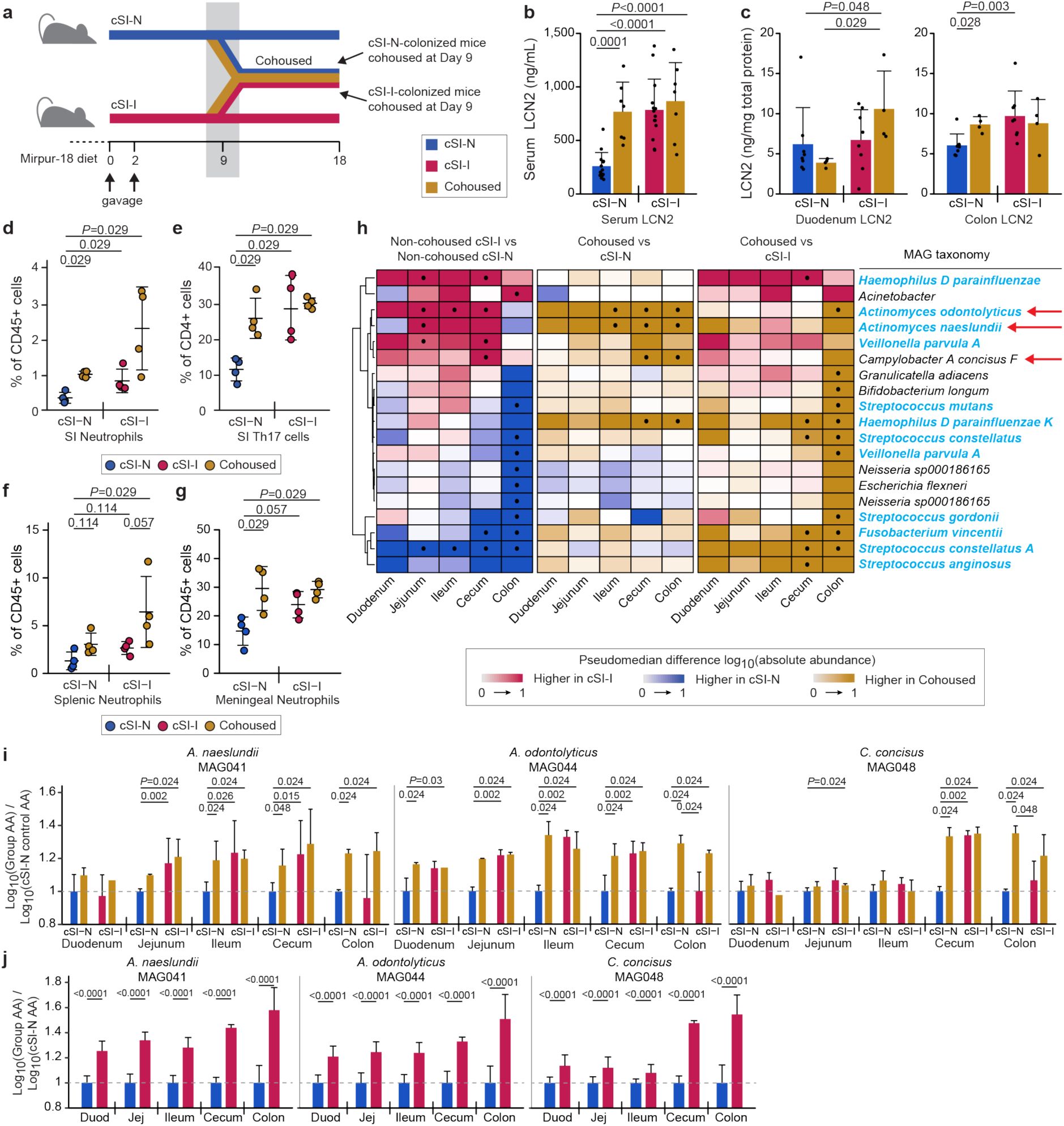
Identification of bacteria associated with pathology after cohousing. **a,** Experimental design of cohousing experiment. **b,** Serum levels of LCN2 protein on experimental day 18, 9 days after initiation of cohousing (n=14 mice/control group, n=7 mice/cohoused group). **c,** Intestinal tissue levels LCN2 measured on experimental day 18, 9 days after cohousing (n=8 mice/control group, n=4 mice/cohoused group). **d,e,** Frequency of neutrophils (**d**), and Th17 cells (**e**), in the small intestinal lamina propria (n=4 mice/group; duodenal, jejunal and ileal segments were combined prior to analysis). **f,g,** Frequency of neutrophils in the spleen (**f**), and meninges (**g**) (n=4 mice/group). For panels **d-f**, each point represents an individual animal; the color code matches that used in panel **a**. **h,** Differences in absolute abundances of MAGs along the length of the gut (n=6 mice/group). MAGs were included if they demonstrated statistically significant differences in their absolute abundance for any comparison between groups in at least one segment of the intestine. MAG taxonomy colored in blue indicates correspondence to ASVs representing ‘core taxa’ in the duodenal microbiota of stunted Bangladeshi children with EED in the BEED study. The three red arrows point to MAGs defined as ‘pathology-associated’. • *P*-adj < 0.05, FDR-corrected Wilcoxon rank-sum. **i,j,** Log_10_ absolute abundances (AA) of the pathology-associated MAGs (MAG041, MAG044 and MAG048) along the length of the intestine in either the cohousing experiment (**i**) or P37 animals from the intergenerational experiment (**j**). Values are expressed relative to cSI-N non-cohoused controls (n=6 mice/control group and n=3/cohoused groups, **i**) or P37 cSI-N offspring in the intergenerational transmission experiment (n=22-25 mice/group, **j**). For **b-j**, *P-*values were determined by Wilcoxon rank-sum test; mean values ± s.d. are shown in **b-g**, **i-j**.

Analogous to P37 offspring from the intergenerational transmission experiments, non-cohoused cSI-I mice and all cohoused animals harbored significantly higher proportions of neutrophils and Th17 cells and a reduced percentage of Th1 cells in the small intestine lamina propria compared to non-cohoused cSI-N controls; this was the case regardless of the initial colonizing bacterial consortium (**Figs. 3d,e**; **Supplementary Table 2f**). Splenic and meningeal neutrophils were similarly elevated in cSI-I and cohoused animals compared to non-cohoused cSI-N mice. **Figs. 3f,g**; **Supplementary Table 2f**). There was no evidence for a mitigating effect of the cSI-N community on pathology in cSI-I colonized mice (**Supplementary Tables 2a,f**). These results suggested either restructuring of strain composition within the cSI-N consortium after cohousing or transfer of bacteria from cSI-I to cSI-N animals, inducing pathology in these formally healthy animals.

We performed shotgun sequencing of intestinal contents from animals in all treatment groups, seeking taxa (MAGs) that were significantly more abundant in the intestines of mice in the ‘inflammatory’ experimental groups (i.e., non-cohoused cSI-I controls and cohoused mice) compared to their ‘non-inflammatory’ counterparts (non-cohoused cSI-N mice). Three MAGs satisfied these criteria: two belonged to the genus *Actinomyces* (*Actinomyces naeslundii* [MAG041], *Actinomyces odontolyticus* [MAG044]) and the other to the genus *Campylobacter* (*Campylobacter concisus* [MAG048]) (**Fig. 3h**; **Supplementary Table 1d**).

Compared to cSI-N controls, the absolute abundance of *C. concisus* was significantly higher in the cecum and colon of cohoused animals, regardless of whether the cohoused animals were initially colonized with the cSI-N or cSI-I consortia (**Fig. 3i**). The absolute abundances of both *A. naeslundii* and *A. odontolyticus* were significantly higher in the jejunum, ileum, cecum and colon of cohoused animals compared to non-cohoused cSI-N controls **(Fig. 3i**). The abundances of all three MAGs were equivalent in cohoused animals and non-cohoused cSI-I controls along the length of the small intestine and in the cecum (**Fig. 3i**), suggesting infiltration of these taxa from cSI-I to cSI-N animals during cohousing. Moreover, the absolute abundances of these MAGs were significantly increased along the length of the gut of P37 cSI-I compared to cSI-N offspring in our intergenerational transmission model (**Fig. 3j**; **Supplementary Table 1d**). *C. concisus* and *Actinomyces spp.* are resident members of the oral microbiota^40^ and are not typically considered to be pathogens, although some strains have been associated with periodontitis and inflammatory bowel disease^41–43^.

Four isolates in our clonally-arrayed culture collection matched these three MAGs that were elevated in cohoused animals compared to cSI-N controls with an average nucleotide identity (ANI) score of >99%: two isolates matched the *A. naeslundii* MAG041 (Bg041a, Bg041b), one matched *A. odontolyticus* MAG044 (Bg044) and one matched *C. concisus* MAG048 (Bg048) (**Supplementary Table 4a;** see **Supplementary Table 4b** for results of mcSEED-based *in silico* metabolic reconstructions of these pathology-associated MAGs, their corresponding isolates, and other phylogenetically related species^44,44^).

### Testing the role of isolates corresponding to pathology-associated MAGs

We directly tested the capacity of these Bangladeshi *C. concisus, A. odontolyticus* and *A. naeslundii* isolates to induce pathology. To do so, germ-free 4-5-week-old C57Bl/6J mice, consuming the Mirpur-18 diet, were gavaged with the cSI-N consortium (experimental day 0). On days 9 through 11, mice were gavaged once daily with either (i) a pool of all four isolates, (ii) the *C. concisus* isolate (Bg048) alone, (iii) the two *A. naeslundii* isolates (Bg041a, Bg041b), (iv) the *A. odontolyticus* isolate (Bg044), or (v) a pool of both *Actinomyces* species. Groups of mice initially colonized with the cSI-N or cSI-I consortium were subsequently gavaged with uninoculated (blank) culture medium as negative and positive controls, respectively (‘sham’ gavage, **Fig. 4a**).

**Fig. 4.**
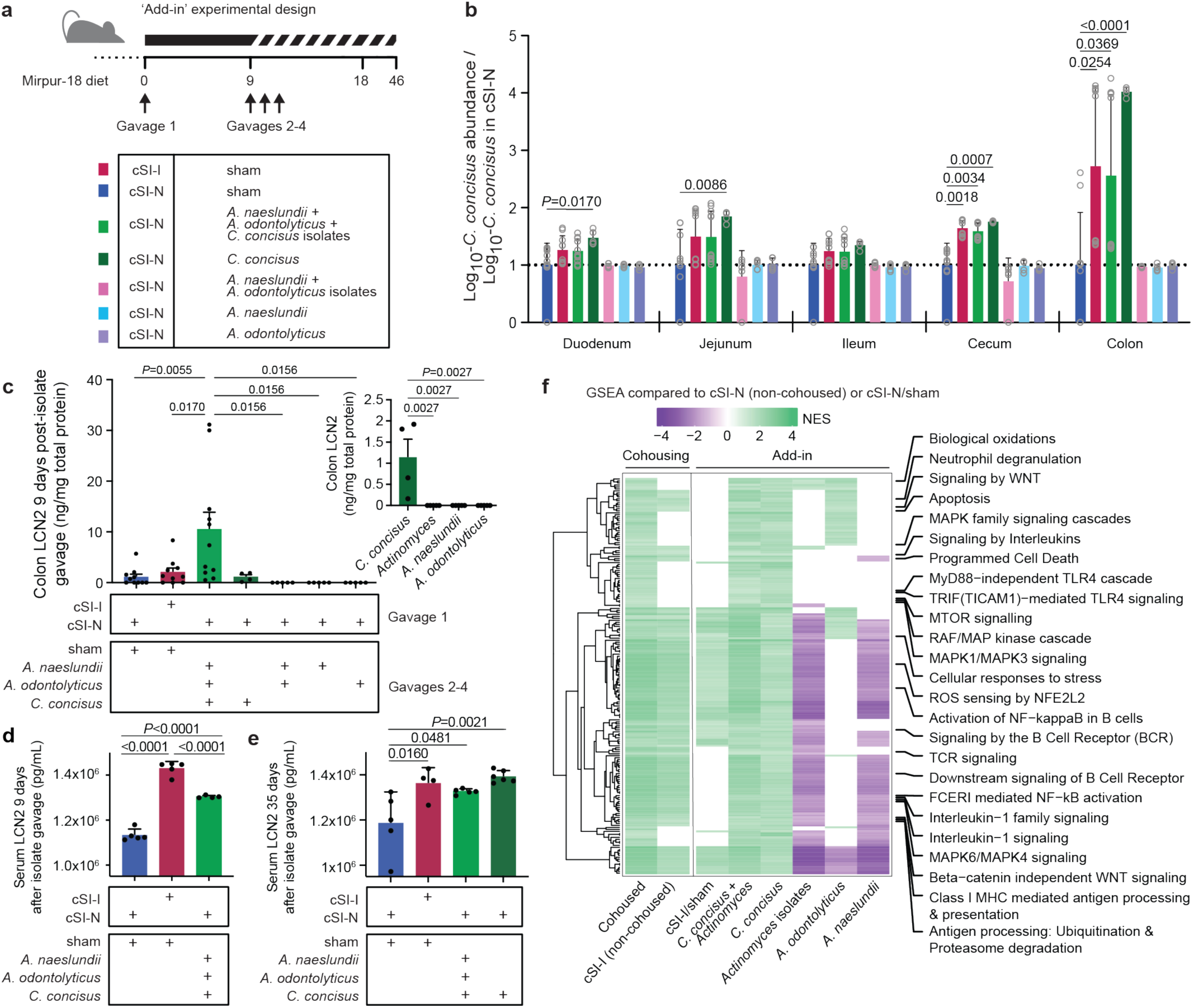
Direct test of candidate pathology-inducing small intestinal bacterial strains in cSI-N mice. **a,** Experimental design of ‘add-in’ experiment. Individual cultured isolates, alone or in combinations, were gavaged once daily on days 9-11 into mice previously colonized with the cSI-N bacterial consortium. **b,** Ratio of the absolute abundance of *C. concisus* strain Bg048 (corresponding to MAG048) shown relative to its absolute abundance in cSI-N/sham controls (a ratio of 1 is indicated by the dotted line; two-way ANOVA with Dunnett’s multiple comparisons to cSI-N/sham control; mean values ± s.d. are shown). Colors correspond to the groups shown in panel **a**. **c,** Levels of LCN2 protein in colonic tissue measured on experimental day 18, 9 days after isolate gavage. The inset shows colonic LCN2 levels after addition of the individual isolates alone (one-way ANOVAs with Tukey’s multiple comparisons; mean values ± s.e.m. are shown). **d,e,** Serum LCN2 at 9 (**c**), or 35 days (**d**), following isolate gavage (mean ± s.d. shown, one-way ANOVAs with Tukey’s multiple comparisons). **f,** Results of bulk-RNA-Seq analysis: normalized enrichment scores (NES) for all Reactome pathways that were significantly enriched in the colon (GSEA *q-*value *<*0.05) of the indicated treatment groups compared to their respective cSI-N counterparts (non-cohoused cSI-N controls, or cSI-N/sham). Pathways related to the immune system are labeled. Pathways were included in the heatmap if they were significantly enriched with the addition of *C. concisus* and in cohoused animals. If a pathway was not significantly enriched, it was assigned a NES score of zero. For panels **b, c, f**, n=5-11 mice/group combined across two independent experiments. For **d** and **e**, n=5 mice/group in a third independent experiment.

Secondary gavage with all isolates combined or with *C. concisus* alone led to an increase in the absolute abundance of *C. concisus*, most notably in the cecum and colon, compared to sham-gavaged cSI-N controls (**Fig. 4b**). The levels of *C. concisus* achieved over the course of the 9 days following its introduction into cSI-N mice were comparable to levels in cSI-I sham-gavaged controls (**Fig. 4b**) as well as in cohoused animals (**Supplementary Table 1d**). In contrast, gavage with the *Actinomyces* species either alone or in concert resulted in a trending, but not statistically significant, increase in absolute abundance along the length of the intestinal tract compared to cSI-N sham-gavaged controls (**Extended Data Fig. 7a,b**; **Supplementary Table 1d**).

Mice colonized with cSI-N then gavaged with *C. concisus* gained significantly less weight than cSI-N sham-gavaged controls or animals gavaged with the *Actinomyces* strains, either alone or together (**Extended Data Fig. 7c,d**). Colonic inflammation, assessed by tissue levels of LCN2, was significantly higher when all isolates (*C. concisus+A. odontolyticus+A. naeslundii*) were gavaged together into cSI-N mice compared to all other treatment groups, including mice receiving *C. concisus* alone and the cSI-I positive controls (**Fig. 4c**). This increase in LCN2 levels did not occur in the duodenum (**Extended Data Fig. 7e**). Similar patterns were observed for the biomarker CHI3L1 (**Extended Data Fig. 7f**). These results suggested that in the distal gut, the presence of *A. odontolyticus* and *A. naeslundii* directly or indirectly affected *C. concisus* in a manner that increased its capacity to induce inflammation.

Systemic inflammation, as quantified by serum LCN2, was significantly higher 9 days after gavage of the pooled strains (*C. concisus+A. odontolyticus+A.naeslundii*) compared to cSI-N controls, but remained lower than in cSI-I mock-gavaged animals (**Fig. 4d**). In a follow-up experiment, serum LCN2 was assessed 35 days after isolate gavage; introduction of either the four isolates together or *C. concisus* alone was sufficient to produce LCN2 levels significantly higher than in cSI-N mock-gavaged controls and not significantly different from cSI-I animals (**Fig. 4e**; **Supplementary Table 2b**). Together, these data suggest that, given a sufficient duration of colonization, this Bangladeshi *C. concisus* isolate can induce levels of systemic inflammation comparable to cSI-I within the context of the cSI-N consortium.

To test whether C. concisus alone could induce intestinal and systemic inflammation, we mono-colonized adult germ-free mice with the organism via oral gavage on experimental days 0, 3, 5, 7, and 14. Two groups served as controls: animals that were orally gavaged with 10^7^ heat-killed *C. concisus* on the same days, and another group that was maintained as germ-free (6-7 mice/group). After 18 days, qPCR assays disclosed 10^2.7^-10^5.8^ organisms/ng DNA in cecal and/or colonic contents in mice gavaged with the live organism; levels were below the limits of detection (10^2^ genome equivalents) in the small intestine of all mice gavaged with live *C. concisus* and for all members of the two control groups (**Supplementary Table 1e**). *C. concisus* levels in the cecal contents of mono-colonized animals were not significantly different from the levels documented in cSI-N mice, and were significantly lower than in animals from the add-in experiment that were first colonized with the cSI-N consortium and then gavaged with *C. concisus* ± the *Actinomyces* strains (118 to 131-fold, respectively), as well as those colonized with the cSI-I consortium (167-fold lower, **Extended Data Fig. 7g, Supplementary Table 1e**). There were no statistically significant differences in duodenal, ileal, colonic, or serum levels of LCN2 nor CHI3L1 between germ-free mice and mice gavaged with heat-killed or live *C. concisus* (P>0.05; repeated measures ANOVA; **Supplementary Table 2b**). These findings indicate that levels of *C. concisus* colonization and its capacity to elicit an inflammatory response are community-context dependent.

#### *C. concisus*-induced inflammatory signaling in the intestinal epithelium

We conducted bulk RNA-Seq of the duodenum and colon to further understand the host response to the presence of *C. concisus*. We compared results across three experimental models in this report: intergenerational dam-to-pup transmission of the cSI-I and cSI-N consortia (P37 offspring), cohousing, and isolate ‘add-in’ experiments (the latter two characterized at experimental day 18) (**Supplementary Table 3d**). We examined Reactome pathways whose expression was significantly enriched (compared to cSI-N controls) in (i) P37 cSI-I animals, (ii) cohoused and non-cohoused cSI-I controls, and (iii) animals initially colonized with the cSI-N consortium subsequently gavaged with *C. concisus*, *A. odontolyticus*, or *A. naeslundii* individually, both *Actinomyces* species, all isolates together, as well as cSI-I/sham controls.

*C. concisus*, either alone or with the *Actinomyces* species, induced a pro-inflammatory transcriptional response in the colon that was not present in the duodenum (**Supplementary Table 3d**). Compared to their cSI-N counterparts, Reactome pathways involved in recruitment and activation of immune cells, reactive oxygen species generation and detoxification, cell death programs, antigen presentation, MAPK signaling, pro-inflammatory cytokines, and toll-like receptor (TLR) signaling were more highly expressed in the colons of P37 cSI-I animals (from the intergenerational transmission experiment), as well as cohoused and non-cohoused cSI-I controls as well as recipients of *C. concisus* (the isolate alone or with *Actinomyces*) (**Fig. 4f**; **Supplementary Table 3d**). Genes involved in cytokine (IL-1) signaling leading to programmed cell death (apoptosis) and immune cell activation (neutrophil degranulation) were induced in the colon following gavage of *C. concisus* alone or in concert with the *Actinomyces* isolates (**Supplementary Table 3d**). Of the 41 immunoinflammatory Reactome pathways that were significantly enriched in the colons of mice gavaged with the three pathology-associated species, only five were enriched in the duodenum (**Supplementary Table 3d)**; none of the 49 Reactome pathways that were enriched in the colon after gavage of *C. concisus* alone were more highly expressed in the duodenum (**Supplementary Table 3d**). The *Actinomyces* isolates, administered alone or together, did not elicit a significant transcriptional response involving Reactome immune pathways, in either the colon or the duodenum (**Fig. 4f, Supplementary Table 3d**). Together, these findings provided evidence that this *C. concisus* isolate, both alone and in concert with the *A. odontolyticus* and *A. naeslundii* strains cultured from the duodenal microbiota of Bangladeshi children with EED, induced an immunoinflammatory gene response that recapitulates aspects of that induced by the full EED donor-derived cSI-I community.

### *C. concisus* metabolism and inflammation

Returning to our previously published V4-16S rRNA dataset generated from the duodenal aspirates of children that took part in the BEED study^2^, we found that the prevalence of *C. concisus* did not exceed the 80% threshold that we had used to define a core set of bacterial taxa in children with EED. Nonetheless, the prevalence of the only *Campylobacter* ASV present (undefined species) was significantly higher in children with ‘epithelial damage’ defined by histopathologic scoring metrics^2^ (**Fig. 5a**, two-sided Fischer’s Exact *P* = 0.0057). The absolute abundance of this *Campylobacter* ASV was significantly positively correlated with levels of only one protein in the duodenal mucosa: CD177, also known as human neutrophil antigen 2 (HNA-2) (Pearson correlation, R = 0.694, *P*-adj = 0.00692). CD177/HNA-2 plays a critical role in neutrophil transmigration and extravasation during an inflammatory response^45,46^.

**Fig. 5.**
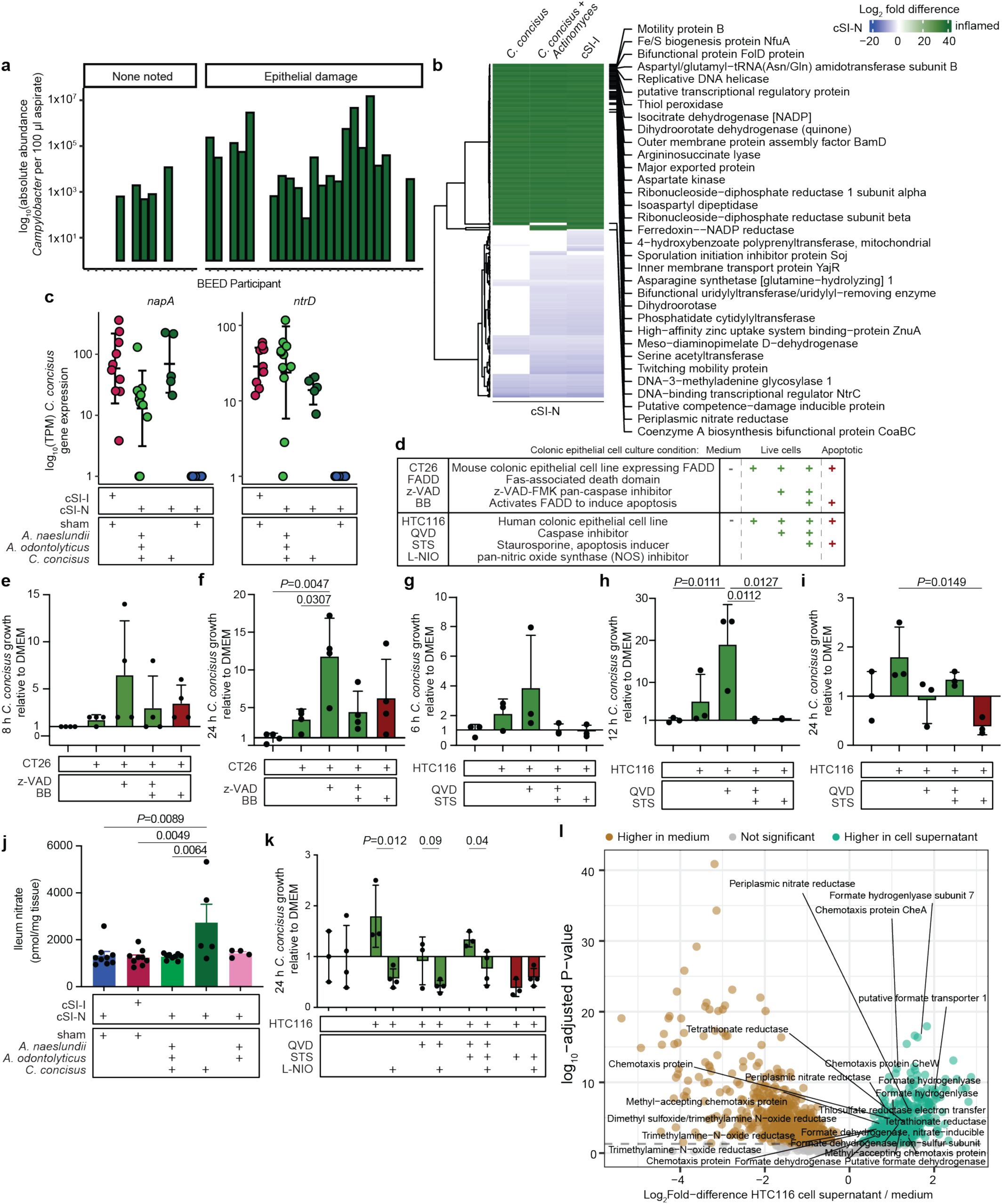
Host epithelial cell nitrate metabolism boosts *C. concisus* growth *in vitro*. **a,** Absolute abundance of the *Campylobacter* ASV identified in duodenal aspirates from children in the BEED study. **b,** Heatmap of significantly differentially expressed genes (*DESeq2* Wald test) in the colon between cSI-I and *C. concisus*-gavaged animals (‘inflamed’) compared to sham-gavaged cSI-N controls. The gene products for the most significantly differentially expressed genes across all three pairwise conditions are labeled. **c,** Expression of *C. concisus* nitrate reductase (*napA*) and nitrate transporter (*ntrD*) 9 days post-isolate gavage, or in cSI-I and cSI-N sham-gavaged controls (tpm, transcripts per million). **d,** Abbreviations of cell lines and treatments used to collect conditioned medium from mouse and human colonic epithelial cells that were live or apoptotic. QVD, Quinoline-Val-Asp-Difluorophenoxymethylketone (caspase inhibitor); L-NIO, *N*^5^-(1-Iminoethyl)-L-ornithine, dihydrochloride (nitric oxide synthase inhibitor). Conditioned medium acquired from live epithelial cells is colored in green; apoptotic cells, in red. **e,f,** Growth of *C. concisus* after 8 (**e**) or 24 hours (**f**) in conditioned medium collected from CT26 mouse colonic epithelial cells that were live (CT26:FADD, z-VAD, z-VAD+BB) or were treated with an inducer of apoptosis (BB). Growth is presented relative to (as a ratio to) growth in cell culture medium (n=4 biological replicates per condition). **g-i,** Growth of *C. concisus* after 6 (**g**), 12 (**h**), or 24 (**i**), hours in spent medium collected from HTC116 human colonic epithelial cells that were live (QVD, QVD+STS), or that had been treated with an inducer of apoptosis (STS) relative to growth in cell culture medium (n=3 biological replicates per condition, growth is presented as a ratio to growth in cell culture medium). **j,** Levels of nitrate in homogenized ileal tissue 9 days post-isolate gavage. **k,** Comparison of *C. concisus* growth after 24 hours in spent medium collected from live (QVD, QVD+STS) or apoptotic (STS) HTC116 cells that were or were not treated with L-NIO (n=3-4 biological replicates per condition, pairwise t-tests for each ± L-NIO comparison). **l,** *C. concisus* genes that were significantly differentially expressed when cultured in conditioned medium collected from HTC116 cells compared to medium alone (*DESeq2* Wald test). For **c-j**, bars denote mean ± s.d.. For **b-c, j**, n=5-11 mice/group combined across two independent experiments. For **e-j**, *P-*values were determined with one-way ANOVA and Tukey’s multiple comparisons.

#### Comparative genomic analysis

Some strains of *C. concisus* decreased expression of tight junction proteins and increased permeability to FITC-dextran in a cultured human adenocarcinoma-derived, enterocyte-like cell line (HT-29)^47,48^. Searches of the Bangladeshi *C. concisus* Bg048 isolate genome and corresponding MAG048 failed to identify known *C. concisus* virulence factors including *Zot* (Zonula occludens toxin), which has been found in 30% of *C. concisus* strains^49^ and reported to disrupt intestinal epithelial tight junctions and augment IL-8 production in HT29 cells^44,47^. Nonetheless, searches within Bg048 and corresponding MAG for homologs to well-described bacterial virulence factors using the Virulence Factor Database (VFDB^50^) identified genes involved in chemotaxis (the chemotaxis regulator *cheY*, flagellar motor switch proteins *fliM* and *fliN*, and flagellum-specific ATP synthase *fliI*) that were conserved with *Campylobacter jejuni* (**Supplementary Table 4a**).

We performed a follow-up comparative genomics analysis that included 119 additional *C. concisus* strains isolated from individuals with oral and/or gastrointestinal pathology (periodontitis, gastroenteritis, Ulcerative Colitis, Crohn’s Disease, Inflammatory Bowel Disease), or healthy controls^51,52^ (**Extended Data Fig. 8**). We performed a functional enrichment analysis that focused on functions (Clusters of Orthologous Genes, COGs) or genes that were enriched in isolates obtained from diseased compared to healthy individuals. COG20 functions and pathways, including genes involved in bacterial defense, bacterial cell wall modification, immune signaling/subversion and Type IV pilus assembly, were significantly enriched in the Bg048 strain and other isolates obtained from diseased compared to healthy hosts (**Supplementary Table 4c**). Our analysis also disclosed that genes encoding a nitrate-inducible formate dehydrogenase (*fdnG*) and formate dehydrogenase H (*fdhF*) were unique to the Bg048 isolate (**Extended Data Fig. 8**). In addition, this strain and only one other among the 119 surveyed contained a *selD* (selenide water dikinase) which, through its capacity to produce selenocysteine, may provide the required selenium cofactor for these formate dehydrogenases^53^ (**Extended Data Fig. 8**, **Supplementary Table 4c**).

#### Intestinal epithelial-derived substrates boost *C. concisus* Bg048 growth

*C. concisus* is known to be non-saccharolytic and have preference for other simple carbon sources (e.g., fumarate, DMSO) and hydrogen gas for growth. Our *in silico* metabolic predictions indicated that *C. concisus* Bg048 has a broad predicted respiratory capacity, including for nitrate, nitrite and nitrous oxide (**Supplementary Table 4b**)^54^.

To garner clues about substrates that support *C. concisus* growth in the gut, we conducted RNA-seq analyses of the cecal microbial community of mice from our ‘add-in’ experiment (**Fig. 4**) and of *C. concisus* grown *in vitro* under anaerobic conditions in rich medium (Bolton Broth with and without 1% mucin). Compared to growth *in vitro* in both media conditions, genes involved in (i) motility (Flagellin A *flaA* and secreted flagellin *flaC*, flagellar ring proteins *flgI/flgH*, assembly factor f*liW*, biosynthetic proteins *fliQ/fliP*, basal body rod protein *flgC*, motility protein *motB*, chemotaxis proteins *cheW*, *cheA*, *mcpH*), (ii) oxidative stress response (superoxide dismutases, *sodB/sodC*), and (iii) several genes encoding respiratory proteins (dimethyl sulfoxide *dorA_2*, *dmsA_1*, *dmsB*, tetrathionate *ttrA/B*, thiosulfate *phsB*, and fumarate *fdrA/B*, *ifcA_2* reductases; formate dehydrogenases *fdhF_3/fdhF_4, fdhB1_2* and hydrogenlyases *hycD/hycG/hycE*) were more highly expressed in cSI-N/*C. concisus*-colonized mice (**Extended Data Fig. 9a**, **Supplementary Table 5a**).

We then compared *C. concisus* gene expression between cSI-I, cSI-N/*C. concisus*, and cSI-N/*C. concisus+Actinomyces* mice compared to cSI-N/mock controls (which harbor lower levels of *C. concisus*) (**Fig. 5b**). The transcript with the largest increase in expression in cSI-N/*C. concisus* mice compared to cSI-N/mock-gavaged animals was a periplasmic nitrate reductase (*napA*, **Figs. 5b,c**). A nitrate import protein (*nrtD*) was also significantly more highly expressed in cSI-N/*C. concisus* mice, suggesting that nitrate may play a role in *C. concisus* virulence (**Fig. 5c**, **Supplementary Table 5b**). Genes involved in (i) motility (*motB*, twitching mobility proteins *pilT_1/pilT_2*, flagellar proteins *flgB, flgG_1/flgG_2, flgH, flgI, flhA, flhB_1/flhB_2, fliG, fliM, fliN, fliP, fliQ, fliR, ylxH*), (ii) redox homeostasis/survival (*sodB*, cytochrome c551 peroxidase *ccpA*, peroxidase *yfeX*, thiol peroxidase *tpx*), and (ii) formate metabolism (*fdhF_1/fdhF_2, hycE*, formate-tetrahydrofolate ligase *fhs*) were also more highly expressed by *C. concisus* in inflammatory conditions (**Fig. 5b, Supplementary Table 5b**).

Having determined that in the context of the cSI-N community *C. concisus* induced an immunoinflammatory response, we tested the hypothesis that *C. concisus* may utilize host substrates that are produced under conditions where there is inflammation and perturbed intestinal epithelial turnover. To do so, we first turned to an *in vitro* tissue culture system in which we cultured mouse- and human-derived colonic epithelial cell lines (CT26 and HTC116, respectively) with or without subsequent induction of cell death (**Fig. 5d**, see *Methods*) and then collected conditioned medium^31^. *C. concisus* growth in the presence of oxygen (5% O_2_) was significantly higher in spent medium from live mouse CT26 cells (**Figs. 5e,f**, **Extended Data Figs. 9b-d**) and human HTC116 cells (**Figs. 5g-i, Extended Data Figs. 9e-g**) compared to conditioned medium harvested from cells undergoing apoptosis or to cell culture medium alone. When cultured anaerobically, *C. concisus* growth did not differ in spent medium from cells that were alive or undergoing apoptosis and was diminished compared to growth in cell culture medium alone (**Extended Data Fig. 9h-k**). These findings suggest that changes in *C. concisus* metabolism in the presence of low levels of oxygen, such as those that may occur in the oxidative state of the inflamed gut^55^, may underlie a fitness benefit conferred to *C. concisus* through utilization of host-derived substrates.

During an inflammatory response, nitric oxide produced by inducible nitric oxide synthase (iNOS, encoded by *Nos2*) reacts with reactive oxygen species to form nitrate^56^. Other Proteobacteria have been shown to utilize host-derived nitrate^56–58^ or other byproducts of iNOS activity^65^ to gain a competitive advantage in mouse models^59^. Given the strong up-regulation of nitrate reductase by *C. concisus* in the gut under inflammatory conditions (cSI-I, cSI-N/*C. concisus*, cSI-N/*C. concisus+Actinomyces*) compared to noninflammatory states (cSI-N mice) (**Figs. 5b,c**), we examined whether *C. concisus* could utilize host-derived substrates generated through metabolism of nitric oxide.

Nitrate in ileal tissue was significantly higher with addition of *C. concisus* alone (cSI-N/*C. concisus*) compared to cSI-N and cSI-I sham-gavaged controls as well as when *Actinomyces* were present (cSI-N/*C. concisus*+*Actinomyces*) (**Fig. 5j**). Nitrate reductase activity reduces nitrate to nitrite; increased bacterial nitrate reductase activity should result in elevated luminal nitrite. Nitrite in cecal contents trended higher in *C. concisus*-gavaged animals (**Extended Data Fig. 9l**), although the difference was not statistically significant, perhaps due to microbial reduction of nitrite or the short half-life of nitric oxides.

To test whether epithelial cell nitric oxide generation contributed to the increase in *C. concisus* growth in spent medium collected from live cells, we treated HTC116 cells with the non-selective NOS inhibitor N-iminoethyl-L-ornithine (L-NIO)^60^. L-NIO treatment significantly reduced the growth benefit conferred to *C. concisus* in conditioned medium collected from live cells when *C. concisus* was grown in microaerophilic conditions. L-NIO had no effect on microaerophilic growth in conditioned medium collected from apoptotic cells (**Fig. 5k, Extended Data Figs. 9m,n**), nor on anaerobic growth in spent medium from live epithelial cells (**Extended Data Figs. 9o,p**). Together, these results suggest that host nitric oxide metabolism boosts growth of *C. concisus* and may confer an advantage under conditions where low levels of oxygen are present in the gut, as is the case with inflammation.

We subsequently profiled *C. concisus* gene expression during growth under microaerophilic conditions in conditioned medium from live HTC116 cell cultures, or the cell culture medium (DMEM). A putative formate transporter (*focA*), formate dehydrogenase (*fdhB*) and formate hydrogen lyases (*hycE, hycD*) plus a periplasmic nitrate reductase *(napA)*, as well as other genes involved in anaerobic respiration were significantly more highly expressed when grown in the presence of live HTC116 spent medium compared to medium alone (**Fig. 5l, Supplementary Table 5c**). Importantly, the periplasmic nitrate reductase was not significantly differently expressed in *C. concisus* grown in conditioned medium from HTC116 cells treated with the NOS inhibitor L-NIO compared to medium alone (DMEM) while other genes involved in anaerobic respiration were still differentially expressed in the presence L-NIO, including those involved in formate metabolism (**Supplementary Table 5c**).

### *C. concisus*-induced inflammatory cytokine signaling enhanced by iNOS

To determine the ‘type’ of immune response elicited by *C. concisus* and whether the host’s capacity to produce nitric oxide plays a role, we collected bone marrow, which harbors a large population of diverse immune cells that seed other body sites, from WT and *Nos2^−/−^* mice and stimulated these cells for 48 hours with live or heat-killed *C. concisus*. ELISA assays established that iNOS was significantly higher after co-culture of wild-type bone marrow cells with live and heat-killed *C. concisus* compared to bacterial medium alone (**Extended Data Fig. 10a,b**). In alignment with our bulk RNA-Seq data of the colon after gavage of *C. concisus* to cSI-N mice (**Fig. 4f)**, IL-1ß secretion was higher in response to heat-killed *C. concisus* and, in a dose-dependent manner, to co-culture with live *C. concisus* compared to medium alone (**Extended Data Fig. 10c,d**). Production of TNF-a (canonical type 1 cytokine), IL-5 (type 2), IL-17 (main cytokine produced by Th17 cells) and IL-22 (type 3 and produced by Th17 cells) were significantly higher after co-culture of wild-type bone marrow cells with live *C. concisus* compared to medium alone (**Extended Data Figs. 10e-j**). In contrast, incubation with heat-killed *C. concisus* elicited a significant increase in IL-5 only (**Extended Data Fig. 10f**). Levels of all five of these cytokines secreted after exposure to live *C. concisus* were significantly lower in bone marrow obtained from *Nos2^−/−^* compared to WT mice. (**Fig. 6a-e**). In *Nos2^−/−^* bone marrow cells, production of IL-1ß, TNF-a, and IL-22 were still significantly higher after incubation with live *C. concisus* compared to the medium control (**Extended Data Fig. 10k-m**). However, genetic ablation of iNOS ameliorated the IL-5 and IL-17 responses – levels in cell supernatants were not significantly higher in response to live *C. concisus* compared to medium control (**Extended Data Fig. 10n,o**). Genes involved in MAPK signaling were more highly expressed in the colon of animals gavaged with *C. concisus* (**Fig. 4f**); p38 MAPK signaling was reduced in *Nos2^−/−^*animals compared to WT, both after exposure to live *C. concisus* and at baseline (**Extended Data Fig. 10p**).

**Fig. 6.**
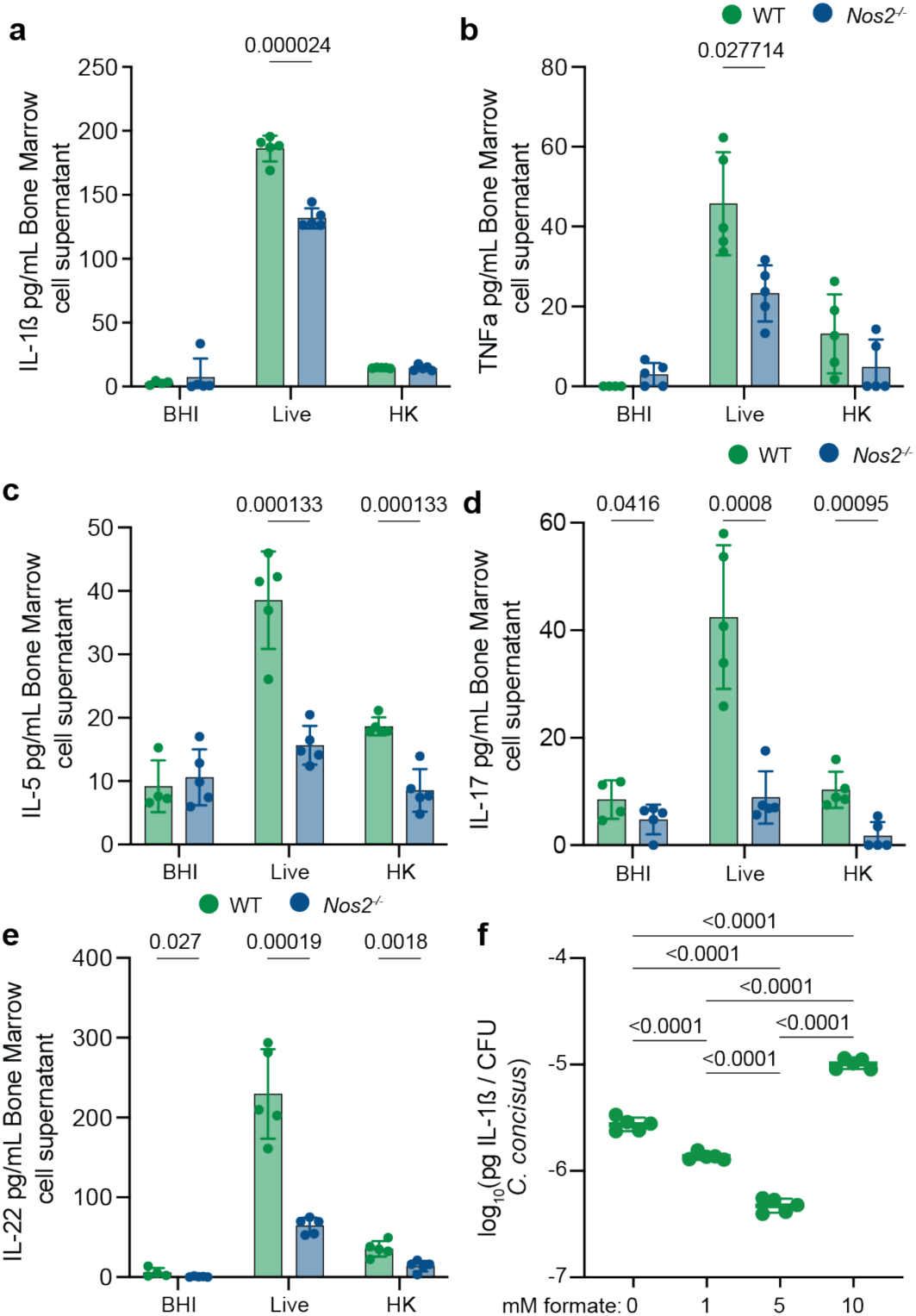
*C. concisus* elicits cytokine signaling in a *Nos2*-dependent manner. **a-e,** Bone marrow cells harvested from conventionally-raised wild-type (WT) or iNOS^−/−^ (*Nos2^−/−^*) mice were incubated with live or heat-killed (HK) *C. concisus* for 48 hours and cytokines were quantified in bone marrow cell supernatants (n=4-5 biological replicates per condition, mean ± s.d., multiple unpaired t-tests with two-stage step-up FDR correction). **f,** IL-1ß quantified in bone marrow supernatants after 24 hours incubation with *C. concisus* pre-treated (grown for the 24 hours prior to co-culture) with various concentrations of formate (n=5 biological replicates per condition, mean ± s.d., one-way ANOVA: F_(3,16)_=527.4, *P*<0.0001, Tukey’s multiple comparisons shown).

As noted above, when *C. concisus* was grown under microaerophilic conditions in spent medium from live HTC116 cell cultures, a nitrate-inducible formate transporter (*fdnG*) and formate transporter (*focA*) were expressed at significantly higher levels (**Extended Data Fig. 10q**).To determine whether formate plays a role in *C. concisus* virulence, as has been described for other pathogens, including *Campylobacter jejuni*^61–63^, we stimulated bone marrow cells with *C. concisus* that had been grown with and without formate. While formate did not increase *C. concisus* growth (**Extended Data Fig. 10r**); (i) live (but not heat killed) *C. concisus* depleted formate when co-cultured with WT or *Nos2^−/−^* bone marrow cells (**Extended Data Fig. 10s**) and (ii) *C. concisus* pretreated with 10 mM formate significantly increased IL-1β produced by bone marrow cells (**Fig. 6f**). Collectively, these findings point to inter-related pathways involving nitric oxide metabolism, formate, and effector cytokines in the host response to *C. concisus*.

## Discussion

Here we describe a preclinical gnotobiotic mouse model of intergenerational transmission of members of the small intestinal microbiota cultured from children with EED that phenocopied several aspects of EED. In the absence of samples from healthy children living in the same environment as those in our clinical study, we generated two bacterial consortia from duodenal aspirates collected from undernourished Bangladeshi children with EED: one encompassing all isolates recovered from the aspirates which induced intestinal and systemic inflammation; the other a subset comprised of one representative strain of each species present in the full collection. This type of ‘reverse translation’ experiment provided an opportunity to perform analyses that were not possible in children in the BEED study who were the source of the small intestinal bacteria, e.g., immune profiling along the length of the gut and in extraintestinal tissues, snRNA-Seq analysis of gene expression in multiple cell lineages in different regions of the small intestine and experimental tests of causality for disease-associated organisms.

We were able to dissect structure/activity relationships in the human donor small intestinal bacterial consortia by combining results from our dam-to-pup intergenerational transmission model with shorter duration cohousing experiments and cultured isolate ‘add-in’ experiments. If the cSI-I and cSI-N consortia had contained unique strains, we would have been able to track the provenance of strains as the two consortia mixed in cohoused mice (and define whether strains ‘bloomed’ or ‘invaded’). However, as the same strains were present in both cSI-I and cSI-N, we are unable to distinguish the origin of strains in cohoused animals. Regardless, from the results of our cohousing experiments, we were able to associate three MAGs – *C. concisus*, *A. naeslundii* and *A. odontolyticus*, all members of the oral microbiota – with development of pathology. Follow-up experiments in which these candidate disease-promoting isolates were introduced to gnotobiotic animals colonized with the non-pathology-inducing cSI-N consortium defined *C. concisus* as a key driver in inducing intestinal inflammation. These experiments will also provide an opportunity for more detailed physiological profiling, including comprehensive energy balance studies, to ascertain the origin of the weight loss phenotype observed in cSI-I mice.

Disruption of the intestinal microbiota by taxa that normally reside in the oral microbiota has been implicated in studies of undernutrition and EED^64,65^. Evidence for this process, referred to as ‘decompartmentalization’ of the microbial community, has come from applying culture-independent methods (primarily16S rRNA amplicon sequencing) to feces. As such, there is little information about disease associations with specific strains, or correlations with their genome-encoded functional features.

*C. concisus* has been reported to be over-represented in the fecal microbiota of stunted compared to non-stunted children living in South India, and significantly more abundant in the feces of a cohort of stunted children living in sub-Saharan Africa compared to healthy controls^66,67^. In children of the BEED study, the only *Campylobacter* ASV present correlated significantly with duodenal mucosal levels of CD177, a cell surface glycoprotein involved in neutrophil activation, and was significantly more prevalent in children with histopathologic evidence of epithelial damage from their mucosal biopsies. *C. concisus* has been described as a typical member of the oral microbiota in healthy individuals (97% prevalence in a cohort of 59 individuals aged 3-80 years)^40^, but it has also been associated with gingival disease^41^ and inflammatory bowel disease (IBD); its prevalence was higher in mucosal biopsies from IBD patients compared to healthy controls^41,42^. The distinctions between *C. concisus* strains recovered from the oral and intestinal microbiota of healthy individuals versus those with immunoinflammatory conditions remain to be fully defined.

It is intriguing that addition of this EED donor-derived *C. concisus* strain to mice harboring the EED duodenal-derived cSI-N consortium of cultured bacteria produced an immunoinflammatory response that was greatest in the colon – at least within the relatively short duration of the cohousing/‘add-in’ experiments which were conducted in coprophagic mice. These findings prompt the question of what factors allow it (and other members of the oral microbiota) to establish themselves in distal regions of the intestine. The isolate ‘add-in’ and mono-colonization experiments described in this report illustrated how inflammation induced by *C. concisus* occurred in a dose- and context-dependent manner; i.e., inflammation in the colon was enhanced in the presence of the *Actinomyces* strains, which themselves were not sufficient to elicit an immunoinflammatory response.

Experiments using heat-killed bacterial culture demonstrated that components and/or products of this pathology-inducing Bangladeshi *C. concisus* isolate were able to induce immune signaling *ex vivo.* Our *in vivo* and *in vitro* studies suggested a role for bacterial nitrate and formate metabolism in pathogenesis of this organism. The mechanisms by which host nitric oxide generation and utilization of host-derived substrates by pathogenic members of the microbiota contribute to decompartmentalization of the oral microbiota and development of enteropathy in humans is an important area that warrants further investigation and could have therapeutic implications. Future studies should also include microbial communities from more distal regions of the intestines of individuals with EED to better characterize how biogeographical features of the microbiota relate to the pathophysiology of this enteropathy.

Our ‘add-in’ experiments indicate that *C. concisus* induces higher levels of inflammation after a longer period of colonization – rather than being cleared by the host. Canonical enteric pathogens, including *Salmonella*^31^, have been shown to utilize substrates from dying, rather than live, intestinal, epithelial cells, as is the case for *C. concisus*. The question of what are the initiating events that allow a pathogen/pathobiont to first establish itself in a microbial community and induce inflammation is complex. One approach is to use genetic tools to identify the contributing signaling and metabolic pathways in both the microbe and host. Our model provides a justification for and sets the stage for these types of detailed analysis.

Endoscopy studies performed as part of the BEED study established that EED is widespread in children as well as undernourished mothers^15,68^; these findings *support* the notion that EED is an intergenerational health problem, at least in some LMICs. Establishing a clinical infrastructure in these areas that allows concomitant sampling of the oral, small intestinal and fecal microbiota of undernourished mothers with EED, as well as healthy mothers without EED, and subsequently creating preclinical models of the type described in this report should provide an opportunity to obtain a more mechanistic understanding of the intricate interplay between the expressed functions of microbial community components, maternal gut health and nutritional status, intestinal adaptations to pregnancy, plus development prior to and following birth. An anticipated outcome of such an effort would be identification of candidate therapeutic targets, including those that may originate in the oral microbiota and reside in the small intestine.

## Acknowledgements

We thank David O’Donnell, Maria Karlsson and the late Justin Serugo for their invaluable assistance with mouse husbandry, Darya Khantakova and Bishan Bhattarai for assisting with flow cytometry preparation and FACS, Martin Meier for generating libraries for shotgun sequencing and microbial RNA-Seq, and Jessica Hoisington Lopez, MariaLynn Crosby and members of the Genome Technology Access Core at Washington University School of Medicine for sequencing these libraries.

## Contributions

Duodenal aspirates used for generating the culture collections were collected in the BEED study, which was directed by T.A.. K.M.P., C.K., M.J.B., and J.I.G. designed the gnotobiotic mouse experiments which were performed by K.M.P., C.K. and R.Y.C.. Histomorphometric analyses were conducted by C.K. and A.E.B.. Flow cytometry and FACS were performed by B.D. and M.L. and interpreted with input from M.C.. ELISA and Luminex assays of serum and tissue proteins were conducted by C.K., R.Y.C., K.M.P., and K.N.. Shotgun sequencing of community DNA, snRNA-Seq and bulk RNA-Seq of intestinal segments, and microbial RNA-Seq data were generated and analyzed by K.M.P., C.K., R.Y.C. and Y.W.. MAG assembly and calculations of MAG abundances were performed by C.K., D.M.W. and M.C.H. Long-read shotgun sequencing and assembly of the genomes of the nominated pathology inducing strains from the culture collection were generated by C.K.. H.L. designed and performed the *C. concisus* qPCR assay. Intestinal epithelial tissue culture experiments were designed by L.K., C.M., and K.R. and conducted by L.K.. *C. concisus* culture and co-culture with immune cell experiments were designed by K.M.P., J.I.G. and M.C., and were conducted and analyzed by K.M.P.D.R. generated *in silico* metabolic reconstructions of MAGs and cultured bacterial strains. K.M.P. performed the comparative genomics analysis of *C. concisus* strains. K.M.P. and J.I.G. wrote the paper with assistance from C.K., M.J.B, and other co-authors.

## Data availability

Datasets generated by (i) shotgun sequencing of DNA isolated from the intestinal contents of gnotobiotic mice and from individual cultured bacterial strains, (ii) snRNA-Seq and bulk RNA-Seq of intestinal tissue, and (iii) microbial RNA-Seq of cecal contents harvested from gnotobiotic mice, have been deposited at the European Nucleotide Archive (ENA; https://www.ebi.ac.uk/ena) under accession number ERA22531095.

## Ethics and Inclusion

This work was performed as part of a long-standing collaboration investigating the role of the gut microbiota in undernutrition that is covered by a memorandum of understanding, between teams led by Tahmeed Ahmed (International Centre for Diarrhoeal Disease Research, (icddr,b) Bangladesh) and Jeffrey Gordon (Washington University in St Louis, USA). Bacterial isolates used in this study were obtained from duodenal aspirates collected from the previously reported Bangladesh EED study, which was approved by the Ethical Review Committee (ERC) at the icddr,b (protocol no: PR-16007; ClinicalTrials.gov number, NCT02812615). Written informed consent was obtained from each child’s parent or guardian to participate and to undergo EGD if the child met the inclusion criteria. Human biospecimens were transferred to Washington University under a Materials Transfer Agreement (MTA). Bacterial isolates cultured and used in this study are the property of icddr,b and are available under MTA upon request to T.A. and J.I.G.

## Code Availability

The custom source code for annotation of bacterial genes and prediction of metabolic pathway presence/absence is available at https://github.com/rodionovdima/PhenotypePredictor-2 and has been accessioned at Zenodo (DOI: 10.5281/zenodo.13250652). All other analyses presented utilize published algorithms with their accompanying methods and are referenced as such.

## METHODS

### Mouse experiments

All experiments involving mice were performed using protocols approved by Washington University Animal Studies Committee. Mice were housed in plastic flexible film gnotobiotic isolators (Class Biologically Clean Ltd., Madison, WI) at 23 °C under a strict 12-hour light cycle (lights on at 0600h). Autoclaved paper ‘shepherd shacks’ were kept in each cage to facilitate the natural nesting behaviors and for environmental enrichment.

#### Source of cSI-I and cSI-N bacterial consortia

The methods used for culturing bacterial strains from duodenal aspirates that were obtained from children enrolled in BEED study are described in a previous publication^5^. Strains were stored at −80 °C in PBS containing 15% glycerol (v/v). For isolates originally isolated under anaerobic conditions, a 20 μL aliquot of each stock was used to inoculate a well in a 1 mL deep-well plate (Thermo Scientific) containing 600 µL LYBHI broth (Brain Heart Infusion [BHI] broth supplemented with 0.05% L-cysteine HCl [w/v] and 0.5% yeast extract [w/v]) followed by incubation under anaerobic conditions (atmosphere 75% N_2,_ 20% CO_2_ 5% H_2_). For isolates originally isolated under microaerophilic conditions (85% N_2_, 10% CO_2_, 5% O_2_), a 20 µL aliquot of each stock was inoculated into 600 µL BHI broth. After incubation for 48 hours at 37 °C, a 20 μL aliquot of each anaerobic or microaerophilic culture was added to 600 μL of fresh medium which was incubated for an additional 24 hours at the same temperature under the same atmospheric conditions. Equal volumes of each isolate sub-culture were then pooled and a solution of PBS containing 30% glycerol (v/v) was added resulting in a final 15% glycerol pooled stock (v/v). Pooling was performed in the Coy chamber. The 15% glycerol pooled stock was sealed in multiple 1.8-mL crimp glass vials (Wheaton) and the vials were stored at −80 °C prior to gavage into mice.

#### Diets

The ‘Adult Mirpur’ diet was designed based on 24-hour dietary recall surveys and food frequency questionnaires taken from adults enrolled in the BEED study living in an urban slum located in the Mirpur district of Dhaka City, Bangladesh^69^. A pelleted, sterile version of this diet was manufactured by Dyets, Inc. (Bethlehem, PA). The quantity of each ingredient used to prepare the diet is provided in **Supplementary Table 1a**. Rice (parboiled, long grain) and red lentils (masoor dal) were each cooked separately with an equal weight of water at 100 °C in a steam-jacketed kettle until the grains were cooked but still firm and then set aside. Tilapia filets (frozen) were steamed separately at 100 °C in a steam-jacketed kettle with a small amount of water until tender (15-20 minutes). Sweet pumpkin (Calabaza variety) was chopped, boiled in the steam-jacketed kettle until soft and then strained. Fresh market white potatoes, Daikon radish (moola), spinach, okra and yellow onions were washed, finely chopped and cooked together in the steam kettle without added water at 70 °C until soft. After cooling, all cooked ingredients were combined and mixed with the wheat flour (atta), soybean oil, salt, turmeric, garlic and coriander powder. The resulting diet was mixed extensively using a planetary mixer, spread on trays, dried overnight at 30 °C, and pelleted by extrusion (½ inch diameter; California Pellet Mill, CL5). Dried pellets were aliquoted into ~250 g portions and placed in a paper bag with an inner wax lining, which was then placed in a plastic bag. Bags were subsequently vacuum-sealed, and their contents sterilized by gamma irradiation (30-50 kGy; Sterigenics).

The composition of the ‘Mirpur-18’ diet was based on Bangladeshi complementary feeding practices for 18-month-old children living in Mirpur, as defined by quantitative 24-hour dietary recall surveys conducted in the MAL-ED study^70^. This diet was manufactured according to a previously described protocol^2^.

The sterility of irradiated diets was confirmed by culture in (i) BHI broth, (ii) Nutrient broth, and (iii) Sabouraud-dextran broth (all from Difco) for one week at 37 °C under aerobic conditions, and in reduced Tryptic Soy broth (Difco) supplemented with 0.05% L-cysteine HCl under anaerobic conditions. All diets were stored at −20 °C prior to use. Nutritional analysis of each irradiated diet was conducted by Nestlé Purina Analytical Laboratories (St. Louis, MO) (**Supplementary Table 1a**).

#### Husbandry for initial tests of bacterial consortia derived from children with EED

4-5 week-old germ-free C57Bl/6J mice were given *ad libitum* access to a standard chow diet (Diet 2018S, Envigo). Three days prior to colonization, mice were switched to the Mirpur-18 diet. For initial comparisons of gnotobiotic mice that had been orally gavaged (oral gavage needle; Cadence Science; catalog no. 7901) with the cSI-I consortium, the cSI-N consortium or the cecal contents of conventionally-raised C57Bl/6J animals that had been maintained on the standard chow diet, one group of animals per treatment arm was euthanized 9 days after gavage, and the second group 28 days later (n=10 mice/group). All animals were weighed and fecal samples were collected three times per week during the experiment.

To test of the role of diet in mediating inflammation, adult female germ-free mice were given breeder chow (Lab Diet 5021) *ad libitum* prior to oral gavage with the cSI-I or cSI-N consortium. Mice maintained on breeder chow for one week prior to switching to the Adult Mirpur diet for the following week and then returned to breeder chow for the final week. Blood was collected from all animals in their isolators retro-orbitally prior to a diet switch (n=3 animals/colonization group euthanized at the end of 7 days on a given diet; the remaining n=5 animals/group were euthanized at 28 days.

#### Husbandry for intergenerational transmission experiments

6-8-week-old germ-free female C57Bl/6J mice were given *ad libitum* access to an autoclaved breeder chow (Lab Diet 5021; Purina Mills, Richmond, IN) until 3 days prior to colonization, at which time they were switched to the Adult Mirpur diet for the remainder of the experiment. Mice received 200 µL of the stock solutions of the cSI-I or cSI-N consortium via an oral gavage needle. Another group of mice received an oral gavage of clarified cecal contents pooled from conventionally raised C57Bl/6J animals maintained on the standard chow (200 μL/recipient animal).

One week after initial gavage, trio matings were performed (two colonized females with one germ-free C57Bl/6J male). Females were continued on the Adult Mirpur diet. Pups born to these mothers were maintained with their dams in the same cage until weaning, at which time pups from the same litter were transferred to new cages and weaned onto the Mirpur-18 diet. For dams and their pups, bedding was replaced every 7 days and diets were provided *ad libitum*. Following weaning of pups on postnatal day 21 (P21), trio matings were performed again using previously pregnant mice. Offspring were euthanized on P37 (first pregnancy) or P14 (subsequent pregnancies) without prior fasting. All biospecimens were flash-frozen in liquid nitrogen and stored at −80 °C until analyses were performed.

For additional matings to assess fetal weights, adult germ-free female C57Bl/6J mice were switched to the Adult Mirpur diet 3 days prior to gavage with 200 µL of the cSI-I or cSI-N consortium, as described above. Two weeks after gavage, female mice were mated in trios and pregnant dams were euthanized at embryonic day (E) 11.5 or E17.5.

#### Husbandry for cohousing experiments

Germ-free 4-5-week-old male C57Bl/6J mice were switched from a standard chow diet (Diet 2018S, Envigo) to the Mirpur-18 diet 3 days prior to colonization. Each mouse received 200 µL of the cSI-I or cSI-N bacterial consortium via a single oral gavage. Nine days later mice were either maintained in their cage and isolator (non-cohoused control groups; n=6-8/group/experiment) or were transferred to a new cage in a new isolator to be cohoused with an equal number of mice transferred from an isolator containing the other experimental group (n=6-8/group/experiment). Bedding was replaced every 7 days for all groups of animals and the Mirpur-18 diet was provided *ad libitum*. All animals were weighed three times a week and euthanized without prior fasting on experimental day 18. All biospecimens were flash-frozen in liquid nitrogen and stored at −80°C before use.

#### Husbandry for tests of candidate mediators of EED (‘add-in’ experiments)

The design of this experiment was analogous to that of the cohousing experiment. Germ-free 4-5-week-old male C57Bl/6J mice were switched from a standard chow diet (Diet 2018S, Envigo) to the Mirpur-18 diet 3 days prior to initial colonization. Each mouse received 200 µL of the cSI-N consortium via a single oral gavage; all mice were maintained in a single gnotobiotic isolator. Nine days later, mice were distributed to new isolators and subsequently gavaged with 200 µL of either 15% PBS/glycerol alone (sham), the cSI-I bacterial consortium, or the cultured strains representing the *A. naeslundii*, *A*. *odontolyticus* and *C. concisus* MAGs. Mice received subsequent 200 µL gavages on days 10 and 11. Another reference control group of mice in their own isolator were initially gavaged with 200 µL of the cSI-I consortium, followed by gavage with medium alone on days 9 through 11. Bedding was replaced every 7 days and the Mirpur-18 diet was provided *ad libitum*. All animals were weighed three times a week and euthanized on experimental day 18 (first round of experiments) or 35 (follow-up more prolonged exposure experiment).

#### Husbandry for mono-colonization experiment

Adult female germ-free C57BL/6J mice were fed the Mirpur-18 diet *ad libitum* starting three days prior to gavage. One group of animals were orally gavaged with (i) 200 µL of a *C. concisus* monoculture grown in X medium (corresponding to 8.7 x 10^6^ CFU) on experimental days 0, 3, 5, 7 and 14. Another group received 200 µL of an equivalent number of cultured cells that had been heat killed (100°C for Y minutes), at the same time points as those used for aministration of live cells. A third group of mice was maintained germ-free. All animals were euthanized on experimental day 18.

#### Division of the intestine

At the time of euthanasia, the small intestine was removed and divided into thirds. Each third was then subdivided into two equal length subsegments (‘proximal’ and ‘distal’). The proximal subsegment was used for either histomorphometric analysis or flow cytometry. Intestinal contents were removed from the distal subsegment by gentle extrusion. The distal subsegment was then cut into three 1.5 cm-long pieces (labeled #1-3 based on their proximal-to-distal location) which were flash-frozen in liquid nitrogen and stored at −80 °C before use. These frozen pieces were used for bulk tissue RNA-Seq (piece #1), snRNA-Seq (piece #2), or ELISA and Luminex-based protein quantification (piece #3). From the proximal end of the colon, three 1.5 cm long pieces (labeled #1-3 based on their proximal-to-distal location) were flash-frozen for the same type of assays applied to small intestinal samples.

#### Histomorphometric and immunocytochemical analyses of intestinal tissue

Proximal subsegments of the duodenum and ileum (see *“Division of the intestine”* above) were fixed in 10% neutral buffered formalin for 24h at 4 °C followed by a 70% ethanol wash. These segments were the placed in parallel with one another and collectively embedded in paraffin. Five μm-thick sections were prepared and stained with hematoxylin and eosin and imaged using a Zeiss Axioscan 7 Slide Scanner. Ten well-oriented crypt-villus units were selected from each intestinal segment and villus height and crypt depth were quantified using *QuPath* (v0.2.3)^71^. Measurements were performed for 4-6 mice per group with the investigator blinded with respect to experimental group.

Sections (5 μm-thick) were prepared from formalin-fixed, paraffin-embedded duodenal and ileal segments. Slides were de-paraffinized and heat-induced antigen retrieval was performed using Trilogy (Millipore Sigma). Slides were blocked in PBS containing 2% BSA, 5% donkey serum and 0.1% Triton-X (antibody buffer) for 30 minutes at room temperature and treated with primary antibodies (Ki67 anti-Rabbit, 1:300, Abcam AB16667; E-Cadherin anti-Mouse, 1:250, DB Sciences 610181) for 90 minutes at room temperature. Following three cycles of washing in PBS, slides were incubated with secondary antibodies (Alexa Fluor Donkey anti-Rabbit 488, 1:300, Invitrogen, catalog no. A32790; Alexa Fluor Donkey anti-Mouse, 1:300, Invitrogen, catalog no. A32731) for 30 minutes at room temperature, followed by a wash in PBS, incubation with DAPI (1:1000 dilution, Thermo Scientific, catalog no. 62248) for 5 minutes, and a final wash in PBS. Slides were mounted with Prolong Gold Antifade Mountant (Invitrogen, catalog no. P36930) and imaged with a Zeiss Axio Scan 7 Brightfield/Fluorescence Slide Scanner. Data were analyzed using *QuPath* (v0.2.3)^71^; as with the histomorphometric analysis, the investigator was blinded with respect to treatment group.

#### Micro-computed tomography (µCT) of bone

The femur was harvested from the left rear leg and cleaned of muscle and connective tissue. Femurs were stored in 10 mL 10% neutral buffered formalin at room temperature for 24 hours. Fixed femurs were then washed in 1X PBS for 15 minutes; this wash step was repeated two more times. Femurs were then subjected to a 30% ethanol wash for 30 minutes, a 50% ethanol wash for 30 minutes and a final 70% ethanol wash for 30 minutes. Femurs were subsequently embedded in 2% agarose and scanned with a μCT 40 desktop cone beam instrument (ScanCO Medical, Brüttisellen, Switzerland). For analyses of cortical bone, 100 slices were taken for each sample in the transverse plane, with a 6 µm voxel size (high resolution); slices began at the midpoint of the femur and extended toward the distal femur. The boundaries and thresholds for bone were drawn manually using μCT 40 software. Volumetric parameters were quantified using software associated with the ScanCO instrument.

#### Protein assays of serum and intestinal tissue

Total intestinal protein was extracted by homogenizing a 1.5 cm-long piece of the duodenum, ileum or colon (piece #3, see *“Division of the intestine”* above) in 600 μL of a solution of ice-cold T-PER Buffer (Thermo Scientific) with Complete Ultra protease inhibitor (Roche), using Lysis Matrix F beads (MP Bio). The homogenate was centrifuged at 13,000 x *g* for 5 minutes at 4 °C. Total protein concentration in the resulting supernatants was quantified using Micro BCA Protein Assay Kit (Thermo Scientific). Protein concentrations were normalized to 1 mg/mL in PBS with Complete Ultra protease inhibitor (Roche). Intestinal and/or serum levels of LCN2 and IGF-1 were quantified using Mouse Lipocalin-2/NGAL DuoSet ELISA and Mouse/Rat IGF-I/IGF-1 DuoSet ELISA (R&D Systems), respectively, following the manufacturer’s instructions. Intestinal and/or serum levels of several other proteins were measured using a custom Mouse Pre-Mixed Multi-Analyte Kit (R&D Systems; CHI3L1, S100A9, MMP8, CXCL1, IL-17A, IL-17E) and the MILLIPLEX MAP Mouse Bone Magnetic Bead Panel (MilliporeSigma; DKK-1, FGF-23, OPG). Assays were performed on a FlexMap3D instrument (Luminex).

### Flow cytometry of immune cell populations

#### Intergenerational transmission experiments

Subsegments of duodenum, ileum and/or colon (*See “Division of the intestine” above*) were digested and myeloid and lymphoid cells were collected according to methods previously described^5^. Briefly, each subsegment was immediately flushed with cold PBS after dissection to remove luminal contents. Each subsegment was then opened lengthwise and gently agitated for 20 minutes at room temperature in Hanks Balanced Salt Solution (HBSS) supplemented with 15mM HEPES, 10% bovine calf serum (BCS) and 5mM EDTA. Each sample was vortexed and the suspended cells were collected; the remaining tissue fragments were subjected to a second round of gentle agitation and vortexing. The tissue remaining after the second collection was rinsed with cold 1X HBSS prior to digestion with Collagenase IV (Sigma) in complete RPMI-1640 for 40 minutes at 37 °C with gentle agitation. Digests were filtered through 100 μm mesh strainer (Falcon) and subjected to density gradient centrifugation using 40% and 70% Percol solutions (GE Healthcare).

To dissect meninges, skin and muscle overlying the skull as well as the mandibles and bone rostral to maxillae were removed. The remaining skull was placed in Iscove’s Modified Dulbecco’s Medium (IMDM, Sigma Aldrich). The meninges were removed from the skull cap using fine forceps and visualization under a light microscope. Meninges were digested for 20 minutes at 37 °C with 1.4 U/mL of Collagenase VIII (Sigma Aldrich) and 35 U/mL of DNAse I (Sigma Aldrich) in IMDM. Following digestion, the tissue was gently pressed through a 70 μm mesh cell strainer (Falcon). The flow-through material was centrifuged at 450 x *g* at 4 °C for 4 minutes. Spleens were processed in a manner similar to what was used for the meninges with the exception that we performed an additional lysis step with ammonium-chloride-potassium (ACK) Lysis Buffer (Quality Biological) prior to staining.

Cells collected from each sample type were resuspended in ice-cold FACS buffer (2 mM EDTA, 25 mM HEPES, 1% BSA in 1X PBS) and stained for extracellular markers at 1:300 dilution. Dead cells were excluded using Zombie NIR fixable Viability kit. Cells were analyzed on a Cytek Aurora (Cytek Biosciences) and the results using FlowJo (v10.8.1). The fluorophore-labeled monoclonal antibodies used for flow cytometry are listed in **Supplementary Table 6** and gating strategies are presented in **Supplementary Information**.

#### Cohousing and add-in experiments

Mice were injected retro-orbitally (under isoflurane anesthesia 2 minutes before euthanasia) with 2 μg of phycoerythrin (PE)–conjugated anti-CD45 antibody (Biolegend, clone 30-F11) to label intravascular leukocytes. Blood was collected from the retroorbital sinus following euthanasia, centrifuged and lysed using ACK lysis buffer (Quality Biological) for 1 minute at room temperature, followed by addition of 2 mL 1X PBS to inactivate ACK lysis buffer. This lysis step was repeated two more times. The resulting material was subjected to centrifugation at 420 x *g* for 4 minutes. Cell pellets were resuspended in FACS buffer (PBS with 2% bovine serum albumin) and stained for extracellular markers at 1:300 dilution.

Immune cells from intestines, spleen and meninges were collected as described above for the intergenerational experiment. Dead cells were excluded using Zombie NIR fixable Viability kit. Cells were analyzed on a Cytek Aurora (Cytek Biosciences) and the data assessed using FlowJo (v10.8.1). The fluorophore-labeled monoclonal antibodies used for flow cytometry are listed in **Supplementary Table 6**.

### Metagenome-assembled genomes (MAGs)

#### Identification of MAGs

DNA was extracted from flash-frozen cecal contents obtained from P37 cSI-I and cSI-N mice and their dams in the intergenerational experiment, as well as from non-cohoused and cohoused mice in the cohousing experiment. Cecal contents were subjected to bead beating and phenol-chloroform extraction to obtain crude genomic DNA, which was subsequently purified (QIAquick 96 PCR Purification Kit) and quantified (Qubit). The final DNA fragment size distribution was determined using an Agilent Technologies 4200 TapeStation.

Fragmented genomic DNA (400-1000 ng) was prepared for long-read sequencing using a SMRTbell Express Template Prep Kit 2.0 (Pacific Biosciences) adapted to a deep 96-well plate (Fisher Scientific) format. All DNA handling and transfer steps were performed with wide-bore, genomic DNA pipette tips (ART). Barcoded adapters were ligated to A-tailed fragments (overnight incubation at 20 °C) and damaged or partial SMRTbell templates were subsequently removed (SMRTbell Enzyme Cleanup Kit). High molecular weight templates were purified (the volume of added undiluted AMPure beads used was 0.45 times the volume of the DNA solution). A second round of size selection was performed by diluting AMPure beads to a final concentration of 40% (v/v) with SMRTbell elution buffer, after which the resulting mixture was added at 2.2 times the volume of the pooled libraries. DNA was eluted from the AMPure beads with 12 µL of SMRTbell elution buffer. Pooled libraries were quantified (Qubit) and their size distribution was assessed using a TapeStation (Agilent). Libraries were then sequenced to a depth of 1.85 ± 1.5 x 10^9^ reads/sample (mean ± SD) using a Sequel II System instrument (Sequel Binding Kit 3.0 and Sequencing Primer v4, Pacific Biosystems). The resulting reads were demultiplexed and Q20 circular consensus sequencing (CCS) reads were generated (Cromwell workflow conFig.d in SMRT Link). CCS reads were assembled into contigs using *metaFlye* (v2.8.1;^72^) with hifi-error set to 0.003 and other options set to default. Contig quality was evaluated using *checkm* (v1.0.7;^73^). Any contig demonstrating *checkm* ‘Completeness’ ≥85% and ‘Contamination’ ≤ 5% was nominated as a ‘high quality’ MAG. Remaining contigs were subjected to further MAG assembly efforts, first by calculating coverage using *CoverM* (v0.6.1, https://github.com/wwood/CoverM) and subsequently MAG reconstruction using *MaxBin* (v2.2.7;^74^). MAGs were cleaned using *MAGpurify* (v2.1.2;^75^) and subsequently evaluated for quality using *checkm*. In an iterative process, candidate MAGs demonstrating *checkm* Completeness ≥ 85% and Contamination ≤ 5% were again nominated as a MAG. High-quality MAGs from both primary and subsequent MAG assembly efforts were dereplicated using *dRep* (v2.3.2;^76^). Final MAG summary statistics were collected with *checkm* and *quast* (v4.5;^77^) (**Supplementary Table 1c**).

#### Taxonomic classification of MAGs

Taxonomic assignments were initially made by employing the Genome Taxonomy Database Toolkit (*GTDB-Tk* v1.5.1;^78^) and corresponding database (release 95; *14*). Phylogenetic trees of MAGs and closely related reference genomes were generated using the Phylogenetic Tree Building service available from the Bacterial and Viral Bioinformatics Resource Center (BV-BRC;^79^). This service utilizes the Codon Tree method and universal protein families as homology group and analyzes alignments of these proteins identified in each genome using the program RAxML. The genome trees obtained were visualized via iTOL^80^.

#### Determination of the absolute abundances of MAGs and isolate genomes

The absolute abundances of MAGs were determined using previously described methods with minor modifications^81,82^. In brief, ‘spike-in’ bacterial strains whose genomes are easily differentiated from those of gut bacteria were added to each weighed frozen sample of intestinal contents or feces prior to DNA isolation and preparation of barcoded libraries^83^. For intergenerational transmission and cohousing experiments, *Alicyclobacillus acidiphilus* spike-in was added (2.22 x 10^8^ cells/mL suspension; DSM 14558; GenBank assembly accession: GCA_001544355.1)^82^. For isolate add-in experiments, a commercial 1:1 mix of *Imtechella halotolerans* and *Allobacillus halotolerans* was added (2 x 10^7^ cells each per 20 µL suspension; Zymo product numbers LMG 26483 and LMG 24826, respectively).

Intestinal contents and fecal samples were subjected to bead beating and phenol-chloroform extraction to obtain crude genomic DNA which was subsequently purified (QIAquick 96 PCR Purification Kit). Shotgun sequencing libraries were constructed using Nextera XT DNA Library Prep Kit and sequenced on an Illumina NovaSeq 6000 instrument [150 nt paired-end reads; 6.17 x 10^6^ ± 3.75 x 10^7^ reads/sample (mean ± SD) from the intergenerational transmission experiment, 3.10 x 10^6^ ± 7.94 x 10^6^ reads/sample from the cohousing experiments, and 3.03 x 10^7^ ± 1.30 x 10^8^ reads/sample from the isolate add-in experiment]. Sample metadata are listed in **Supplementary Table 7**. Due to increased host reads in small intestinal samples, libraries were balanced based on a host:microbe read ratio determined by preliminary shallow sequencing.

Reads were demultiplexed (*bcl2fastq*), trimmed (*trimgalore,* v0.6.1; https://github.com/FelixKrueger/TrimGalore) and filtered to exclude host reads (*bowtie2*; v2.3.5;^84^). MAG abundances were determined by assigning reads to each MAG, followed by a normalization for genome uniqueness in the context of a given community^83^. The resulting counts table was imported into R (v4.0.4). We calculated the absolute abundance of a given MAG *i* in sample *j* using the following equation:

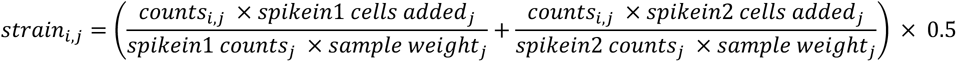

The statistical significance of observed differences in the abundances of a given MAG across different groups was tested using a non-parametric Wilcoxon-rank test with FDR corrections on calculated log_10_ absolute abundances. Final MAG abundances for all experiments are included in **Supplementary Table 1d**.

#### Identification and assembly of isolates genomes related to MAGs

Average nucleotide identity was calculated using *dRep* (v2.3.2;^76^) for all genomes assembled by shotgun sequencing (paired end 150 bp reads) of cultured isolates from the BEED study duodenal aspirates (**Supplementary Table 1b**)^2^. For cultured isolates displaying ANI ≥ 95% shared with MAGs of interest obtained from the intergenerational and cohousing experiments, monocultures of each isolate were grown in 10 mL broth overnight at 37 °C under anaerobic conditions (atmosphere; 75% N_2_, 20% CO_2_ and 5% H_2_) without shaking. Cells were recovered by centrifugation (5,000 x *g* for 10 minutes at 4 °C) and high molecular weight genomic DNA was purified (QIAquick 96 PCR Purification Kit) and quantified (Qubit); the final fragment size distribution was determined using a TapeStation (Agilent). Fragmented genomic DNA (400-1000 ng) was prepared and long-read sequencing was performed as described above. The resulting reads were demultiplexed and Q20 circular consensus sequencing (CCS) reads were generated (Cromwell workflow conFig.d in SMRT Link). Genomes were assembled using *Flye* (v2.9;^85^) with hifi-error set to 0.003 and other options set to default. Assembly quality was evaluated using *checkm*. Taxonomic assignments were initially made by employing the Genome Taxonomy Database Toolkit (*GTDB-Tk* v1.5.1;^78^) and corresponding database (release 95; *14*). Final assembly summary statistics were collected with *checkM*, *quast* and *dRep* (**Supplementary Table 4a**). All isolates were of high quality based on marker gene analysis, consisted of one or two contigs, and shared >99.9999% ANI with their respective MAG (**Supplementary Table 4a**).

#### In silico metabolic reconstructions of MAGs and isolate genomes

MAGs and isolate genomes were initially annotated using *prokka* (v1.14;^86^); KEGG Orthology (KO) numbers were assigned to bacterial coding sequences using BlastKOALA^87^. MAGs were then subjected to *in silico* metabolic pathway reconstruction. Our approach for reconstruction is based on gene annotation using the microbial community SEED (mcSEED) database containing 2,856 reference bacterial genomes for which metabolic phenotypes have been inferred from expert curation and subsequent *in silico* predictions of the presence or absence of the pathway in each genome^88,89^. These results capture utilization and/or biosynthesis of 106 metabolites, including carbohydrates, amino acids, vitamins, and fermentation end-products^90–92^.

For functional annotation of coding sequences in each genome, we used a combination of public domain tools (such as DBSCAN, DIAMOND, MMSeq2), custom scripts and the mcSEED reference database genes. These annotations were supplied to a ‘Phenotype Predictor’ pipeline that allows automated propagation of curated metabolic phenotypes over new microbial genomes and MAGs from represented phylogenetic groups using a consensus of three complementary approaches: (i) rule-based phenotype assignment using the genomic distribution of orthologs and a set of phenotype rules; (ii) machine learning models for binary phenotype prediction trained on reference sets of genes and phenotypes and (iii) a neighbor-based approach for phenotype assignment based on variability of metabolic phenotypes and genes within a group of phylogenetically close neighbors. The combination of all three approaches allows consensus phenotype assignment with >99% accuracy^93^. The mcSEED gene annotations are provided in **Supplementary Table 4b**.

### Quantitative Polymerase Chain Reaction (qPCR)

To determine the absolute abundance of *C. concisus* in mono-colonized animals, we performed qPCR on DNA extracted from duodenal, jejunal, ileal, cecal, and colonic contents. DNA was extracted using the QIAamp Powerfecal Pro DNA kit and amplified in quadruplicate 10 µL reactions that contained 5 µL iTaq Universal SYBR Green Supermix (Bio-Rad), 0.4 µL of a 10 µM stock solution of primers targeting the *C. concisus cpn60* gene^94^ (JH0023: GGCTCAAAAGAGATCGCTCA; JH0024: CCCTCAACAACGCTTAGCTC), and 1.36 μL of a 3 ng/µL solution of DNA isolated from intestinal contents or 2.5 uL of a 1ng/uL reference control solution of genomic DNA isolated from a monoculture of *C. concisus)*. Amplifcation was performed using a QuantStudio 6 Flex machine and the following cycling conditions; 95°C for 3 minutes, followed by 40 cycles of 15 seconds at 95°C, 15 seconds at 64.6°C, 15 seconds at 72°C, and a final melt at 95°C for 1 minutes. *C. concisus* genome equivalents in DNA extracted from mouse intestinal samples was calculated using standard curve of Ct values generated from the purified preparation of *C. concisus* genomic DNA.

### Bulk RNA-Seq

#### Intestinal tissue

RNA was extracted from 1.5 cm flash-frozen duodenum, ileum and/or colon (piece #1, *See “Division of the intestine” above*) using the Qiagen RNeasy 96 Kit. Total RNA was quantified (Qubit) and quality was assessed using a TapeStation (Agilent). cDNA libraries were generated using the Illumina Total RNA Prep with Ribo-Zero kit (Illumina). Barcoded libraries were sequenced on Illumina NovaSeq 6000 instrument [150 nt paired-end reads to a depth of 3.23 x 10^7^ ± 3.54 x 10^6^ reads/sample (mean ± SD) from the intergenerational experiment and 7.13 ± 3.08 x 10^7^ reads/sample from the cohousing experiment)]. Sample metadata are provided in **Supplementary Table 7**.

#### Data analysis

Read quality was verified with *FastQC* (v0.11.7; http://www.bioinformatics.babraham.ac.uk/projects/fastqc/). Reads were then trimmed to remove adapters and low-quality read segments using *trimgalore* (v0.6.1). Trimmed reads were then pseudoaligned to a *kallisto* (v0.46.2^95^) index built from the Gencode v25 *Mus musculus* reference genome. Transcript counts were aggregated to gene counts. *Kallisto* ‘estimated counts’ values and transcripts per million (TPM) values were used to generate a gene-level expression matrix in R using *tximport* (v1.22.0;^96^) and *biomaRt* (v2.50.3;^97^). The ‘lengthscaledTPM’ function in *tximport* was used to adjust estimated counts by gene length and abundance. This resulting count matrix was imported into *limma-voom* (v3.50.3;^98,99^) to compare log_2_-cpm gene expression between groups. GSEA was performed using *clusterProfiler* (v4.2.2;^100^) and the Gene Ontology (GO) pathway database, with FDR-correction.

### snRNA-Seq

#### Isolation of nuclei from small intestinal tissue

Nuclei were extracted from flash-frozen 1.5 cm pieces of duodenum and ileum (piece #2, see *“Division of the intestine”* above) collected from P37 mice in the intergenerational transmission experiment. Briefly, the tissue piece was thawed and minced in lysis buffer (25 mM citric acid, 0.25 M sucrose, 0.1% NP-40, 1X protease inhibitor). Nuclei were released from cells using a Dounce homogenizer (Wheaton), then washed 3 times with buffer [25 mM citric acid, 0.25 M sucrose, 1X protease inhibitor (Roche)] and filtered successively through 100 μm-, 70 μm-, 40 μm-, 20 μm- and 5 μm-diameter strainers (pluriSelect) to obtain single nuclei. Following filtration, nuclei were pelleted by centrifugation (500 x *g* for 5 minutes) and resuspended in a buffer containing 5 mM KCl, 3 mM MgCl_2_, 50 mM Tris, 1 mM DTT, 0.4 U/μL RNase inhibitor (Sigma), plus 0.4 U/μL Superase inhibitor (ThermoFisher)^101^.

#### Generation and sequencing of snRNA-Seq libraries

We used approximately 7,000 nuclei per intestinal sample for gel bead-in-emulsion (GEM) generation. Reverse transcription and library construction were performed according to the protocol provided in the 3’ gene expression v3.1 kit manual (10X Genomics PN-1000121). Balanced libraries were sequenced on Illumina NovaSeq 6000 instrument [150 nt paired-end reads; 2.95 x 10^8^ ± 2.02 x 10^7^ reads/sample (mean ± SD)]. Sample metadata are listed in **Supplementary Table 7**.

### Analysis of snRNA-Seq datasets

#### Preprocessing and quality control

Read alignment, feature-barcode matrices and quality controls were performed using the *CellRanger* 5.0 pipeline with the flag ‘--include-introns’ (GRCm38/mm10). ‘Ambient’ RNA was removed using the *remove-background* module from CellBender^102^. Sample integration, count normalization, cell clustering and marker gene identification were performed using Seurat 4.0^103^. Briefly, filtered feature-barcode matrices from CellRanger were imported as a Seurat object using *CreateSeuratObject*. Matrices were filtered to remove low quality nuclei (defined as nuclei with < 200 or > 5000 genes or < 400 UMIs). For intestinal samples, nuclei with over 5% reads from mitochondrial genes or over 5% reads from ribosomal protein genes were excluded. Each sample was then normalized using *SCTransform*^104,105^ and predicted doublets were removed using *DoubletFinder*^106^. Samples of the same tissue type (duodenum or ileum) were integrated using *SelectIntegrationFeatures*, *PrepSCTIntegration*, *FindIntegrationAnchors* and *IntegrateData* from the Seurat software package. Each integrated dataset was subjected to unsupervised clustering using *FindNeighbors* (dimensions = 1:30) and *FindClusters* (resolution = 0.8 and 1.2 for duodenum and ileum, respectively) from the Seurat package.

#### Cell type annotation

Cell type annotation for duodenal and ileal snRNA-Seq objects was performed using the *FindMarkers* function in Seurat. Manual cell type assignments were conducted based on expression of reported markers^107^.

#### Pseudo-bulk analysis of differential gene expression

Genes defined as having low levels of expression (read count < 4) were filtered out prior to count aggregation across nuclei for a given cell type (cluster) within each biological sample. Each pseudo-bulked sample served as input for *DESeq2*-based differential gene expression analysis (likelihood ratio test, minimum=1e-6^108^). Genes with differential expression (adjusted *P* < 0.05) were used as input for GSEA with *clusterProfiler*^100^.

#### Intercellular signaling

Signaling within each tissue (duodenum or ileum) was inferred with *‘Differential’ NicheNet*^35^. Briefly, log-transformed counts were used for as input for the wrapper function *nichenet_seratobj_aggregate*. Default ligands, receptors and target matrices were used. In the duodenum and ileum, all cell clusters were designated as ‘sender’ cells to stem and transit amplifying (TA) ‘receiver’ cells^36^. *Differential NicheNet* includes differential expression of ligands within a given ‘sender’ cell type between conditions (cSI-I vs. cSI-N) in the ranking of ligand-receptor pairs, while the *NicheNet* algorithm does not.

### *C. concisus* comparative genomics analysis

Published *C. concisus* whole genomes (97 from refs^51^, 14 from ref^52^, 8 from NCBI taxid=199) were downloaded from NCBI through *anvi’o* (v8, ref^109^). A typical pan-genomic workflow was conducted. Briefly, whole genomes (.gbff files) and associated metadata were downloaded through their NCBI BioProject accessions or taxid numbers. Contig databases were generated with default config settings using the anvi’o snakemake workflow^110^; NCBI-PGAP was used as the gene caller to create a genomes storage database. For the pangenome analysis^111^, ncbi-blast was used with *mcl-inflation*=10 and *minbit*=0.5. Genome similarity was computed with *pyANI*. Disease-associated metadata were manually curated and used to compute functional enrichment (*anvi-compute-functional-enrichment-in-pan*^110^. The statistical approach used to define enrichment scores for functions or genes fits a generalized linear model to the occurrence of each module and computes a Rao test statistic with multiple hypothesis correction; this approach has been described previously^110^.

### Epithelial cell death induction assay

HCT116 cells were cultured at 37 °C in DMEM medium supplemented with 10% FBS at 4 x 10^5^ cells per ml. Cells were washed with 1xPBS and selected controls were pre-treated with DMSO (0.1%) or 30 μM caspase inhibitor (QVD, Quinoline-Val-Asp-Difluorophenoxymethylketone) for 1h before inducing apoptosis with administration of 1 μM staurosporine (STS; Abcam) for 24 h.

CT26:FADD cells were seeded at 2 x 10^6^ cells per 10 cm culture dish. The next day, cultures were incubated with 1 μg/ml doxycycline for 16 h to induce expression of the FADD construct (Fas-associated death domain). Doxycycline was removed by washing with 1x PBS before 1 h pre-treatment with 20 μM z-VAD-FMK (z-VAD, pan-caspase inhibitor, MedChem Express) or DMSO (0.1%). Cell death was induced by adding 10 nM B/B homodimerizer for 5 h after which time supernatants were collected and centrifuged at 350 x *g* for 5 minutes to remove cellular debris. The resulting supernatants were filtered using 0.2μm syringe filter (SFCA; Corning) and then frozen at −20 °C for later use in *C. concisus* growth assays.

#### *C. concisus* growth in conditioned medium from intestinal epithelial cells

*C. concisus* isolate Bg048 was routinely cultured in Bolton Broth supplemented with 10% FBS (autoclaved and subsequently filter-sterilized) and on BHI-agar plates supplemented with 7.5-10% (v/v) horse or sheep blood. For growth in spent medium collected from colonic epithelial cell lines, 25 µL of a 24-hour culture of *C. concisus* in Bolton Broth with 10% FBS was sub-cultured into a 1:1 (v/v) mixture of Bolton Broth with 10% FBS and intestinal epithelial cell culture spent medium or blank cell culture medium as a control (DMEM). After a 6-24 h incubation at 37 °C, a 30 µL aliquot of the resulting culture was removed for serial dilutions and the determination of CFU using blood agar plates incubated under micro-aerophilic conditions (5% O_2_, 10% CO_2_, 85% N_2_) or anaerobic conditions (5% H_2_, 20% CO_2_, 75% N_2_). After 48 hours of growth, *C. concisus* colonies were counted and growth relative to DMEM was evaluated.

### Microbial RNA-Seq

#### *C. concisus* gene expression in the mouse gut

RNA was isolated from cecal contents harvested from cSI-I and cSI-N sham-gavaged controls, and ‘cSI-N plus isolate-add in’ mice described in **Fig. 4** and **Extended Data Fig. 7**. cDNA libraries were generated from isolated RNA samples using the Total RNA Prep with Ribo-Zero Plus kit (Illumina). Barcoded libraries were sequenced on an Illumina NovaSeq 6000 instrument [150 nt paired-end reads to a depth of 1.93 x 10^8^ ± 7.94 x 10^7^ reads/sample (mean ± SD); n=6 samples/treatment group]. Sample metadata are listed in **Supplementary Table 7**. Raw reads were trimmed (*trimgalore*; v0.6.1), filtered to exclude host reads (*bowtie2*; v2.3.5), and mapped to MAGs (*kallisto*; v0.46.2) and the *C. concisus* isolate genome (see section of *Methods* titled ***Identification and assembly of isolates genomes related to MAGs*)**. The resulting *kallisto* pseudocounts tables were imported into R (v4.0.4). The geometric mean of absolute abundance (see ***Determination of the absolute abundances of MAGs and isolate genomes*)** was used to set *sizeFactors* for the *DESeq2* object prior to differential expression with the *DESeq2 Wald* test.

#### *C. concisus* gene expression in vitro

*C. concisus* was grown anaerobically on solid BHI agar supplemented with 10% sheep blood. After 48 hours, single colonies were picked into rich medium (Bolton broth supplemented with 10% FBS); after 24 hours of anaerobic growth at 37 °C, an aliquot was diluted 1:32 into media used for assessing gene expression; these conditions included (i) Bolton broth supplemented with 10% FBS ± 1% mucin, grown anaerobically and (ii) Bolton broth supplemented with 10% FBS mixed 1:1 (v/v) with conditioned medium from HTC116 colonic epithelial cells, grown under microaerophilic conditions. The colonic epithelial cell media conditions used were as follows; (i) conditioned medium from HTC116 cells grown in DMEM, (ii) conditioned medium from HTC116 cells treated with L-NIO and (iii) DMEM cell culture medium control. After 24 hours, these broth cultures were transferred to 5 mL snap-top microcentrifuge tubes and centrifuged at 20,000 x *g* for 30 minutes at 10 °C to pellet *C. concisus* cells. Supernatant was removed and cell pellets were frozen at −20 °C. RNA was extracted with the Qiagen RNeasy UCP Micro kit. Briefly, 200 µL of acid-washed glass beads were added to the bacterial pellets on ice with 350 µL buffer RULT containing 2-mercaptoethanol. Bead beating was performed for 5 minutes, and followed by a protocol described in the kit, including on-column DNase treatment. Quality was assessed with RNA High Sensitivity ScreenTape using an Agilent Technologies 4200 TapeStation and quantified with Qubit RNA High Sensitivity. ERCC RNA Spike-In Mix (Invitrogen, 4456740) was added prior to library prep which was performed using the Total RNA Prep with Ribo-Zero Plus kit (Illumina). Barcoded libraries were sequenced [AVITI instrument (Element Biosciences); 150 nt paired-end reads; 1.03 x 10^7^ ± 7.4 x 10^6^ reads/sample (mean±SD)]. Demultiplexed reads were quality-trimmed (*trimgalore*) and mapped to the *C. concisus* isolate genome Bg048 and ERCC transcripts (*kallisto*). Kallisto pseudocounts tables were imported into R, and ERCC counts were extracted to create a *DESeq2* object from which*ESeq2 sizeFactors* were calculated and centered. These *sizeFactors* were used to normalize the *C. concisus DESeq2* object prior to differential expression testing (Wald test).

To facilitate comparison of gene expression between the *in vivo* and *in vitro* experimental conditions, reads from housekeeping genes (*rpoA, gyrB, recA, dnaK, atpA*) were extracted; the geometric mean of these five housekeeping genes was calculated per sample and centered for the *DESeq2 sizeFactors* in a manner analogous to the ERCC spike-in control.

### *C. concisus* co-culture with mouse immune cells

Conventionally raised 8-12 week old female wild-type C57Bl6/J or *Nos2^−/−^*(B6.129P2-*Nos2^tm1Lau^*/J) mice fed standard mouse diet (LabDiet 5010) were euthanized. Femurs were collected and placed into RPMI 1640 supplemented with 2 mM L-glutamine and 10% FBS and kept on ice. Bone marrow was extracted from each femur by centrifugation and resuspended in 1 mL RPMI. Briefly, femurs were placed vertically into a 0.65 mL snap-top tube (Avant 2924) with a puncture hole created by an 18 gauge needle. The tube was placed within a 1.5 mL microcentrifuge tube (Eppendorf) and samples were centrifuged >10,000 x *g* for 15 seconds. Bone marrow cells were resuspended in RPMI and pooled between mice (n=2 animals/genotype per experiment) and 200 µL aliquots were placed into wells of a 24-well tissue culture plate. Live *C. concisus* cultures grown anaerobically in Bolton broth for 24 hours or heat-killed at 100 °C for 2.5 hours 400 µl was added to each well containing bone marrow from WT or iNOS^−/−^ mice. Cells were cultured under an atmosphere of 5% CO_2_ at 37 °C for 48 hours. Tissue culture plates were centrifuged at 500 x *g* to pellet cells and collect cell supernatants, which were aliquoted into PCR plates and stored at −20 °C. Cell pellets were washed with 1 mL PBS, centrifuged at 1000 x *g*, and stored at −20 °C until protein extraction and quantification.

#### Quantification of cytokines with ELISAs

Bone marrow cell supernatants were thawed and cytokines were assayed using 50 µL of undiluted following the manufacturer’s instructions (IL-1B, Abcam 197742; IL-17, Abcam 199081; IL-22, Abcam 223857; IL-5, Abcam 204523; TNFa, Abcam 208348; MAPK signaling, Invitrogen, 85-86195-11). To quantify levels of iNOS in bone marrow cell pellets, protein was extracted using the cell lysis buffer provided in the iNOS ELISA kit (Abcam 253219). The cell extract was diluted 1:3 to quantify protein concentration using the Micro BCA Protein Assay Kit (Thermo Scientific 23235). 50 µL of cell extract was used quantify iNOS following instructions provided with the ELISA kit; final iNOS concentration was normalized to total protein concentration.

---

**Extended Data Fig. 1.**
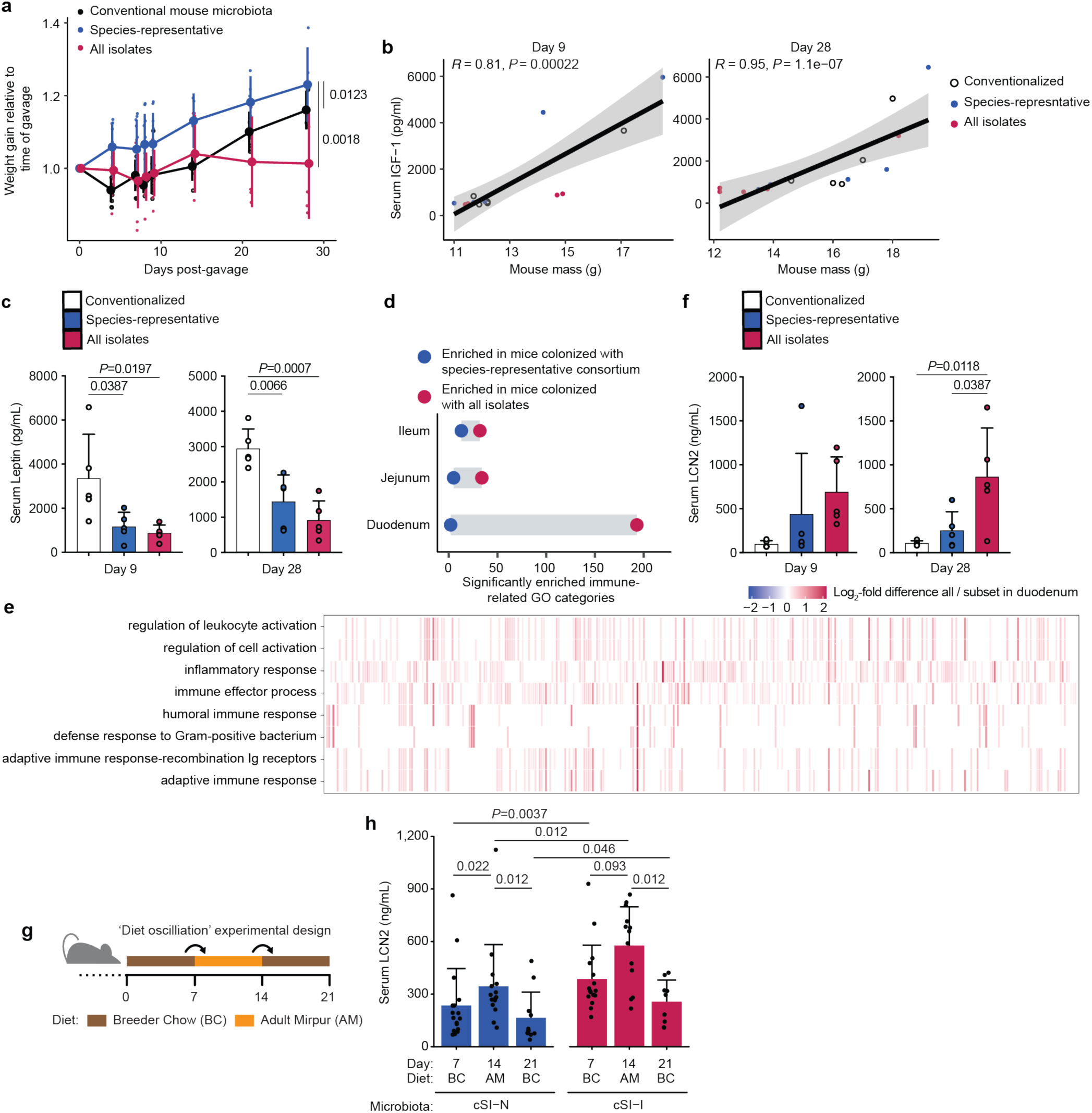
Initial test of the host response to colonization with the cSI-N and cSI-I consortia. **a,** Weight gain of mice colonized with all 184 isolates cultured from duodenal aspirates obtained from children with EED (cSI-I), or a species-representative set of isolates (cSI-N). A control group of CONV-D mice was colonized with cecal microbiota harvested from conventionally raised animals. All mice were fed Mirpur-18 diet *ad libitum*. Each small point represents the weight gain for an individual mouse; larger points denote the mean, error bars, standard deviation. n=5-10 mice/group, linear mixed effects model (Weight ~ Group*Day + (1|MouseID), *P*-values shown are the result of Tukey’s post-hoc tests). **b,** Serum IGF-1 correlates with mouse weight measured 9 (left) or 28 (right) days after colonization (n=5 mice per group per timepoint, linear regression). **c,** Serum leptin levels determined nine (left) or 28 days (right) days post-colonization (n=5 mice per group per timepoint, bars denote mean ± s.d.). **d,** Number of Gene Ontology (GO) categories related to the immune system that were significantly enriched (GSEA *q* value < 0.05) based on differential gene expression in the small intestine of mice colonized with all isolates (red) versus the species representative subset (blue). Bulk RNA-seq was performed in duodenal, jejunal, and ileal tissue 28 days following gavage (n=5 mice/group). **e,** Expression of genes (columns, un-labeled) that comprise the leading edge of the eight GO categories most significantly enriched in animals colonized with the full consortium (red) compared to the species-representative subset (blue). Bulk RNA-seq of duodenal tissue, 28 days post colonization. **f,** Serum lipocalin-2 (LCN2) levels 9 (left) or 28 days (right) days post-gavage of the consortia (n=5 mice per treatment group per timepoint.). **g,** Design of experiment to test whether systemic inflammation induced by cSI-I consortium is influenced by diet. **h,** Serum levels of LCN2 after mice were fed breeder chow for one week (day 7), Adult Mirpur the following seven days (day 14), and a final week of breeder chow (day 21) (Wilcoxon rank-sum tests, n=8-17 mice/group). For **a,c,f,h**, bars denote mean ± s.d.

**Extended Data Fig. 2.**
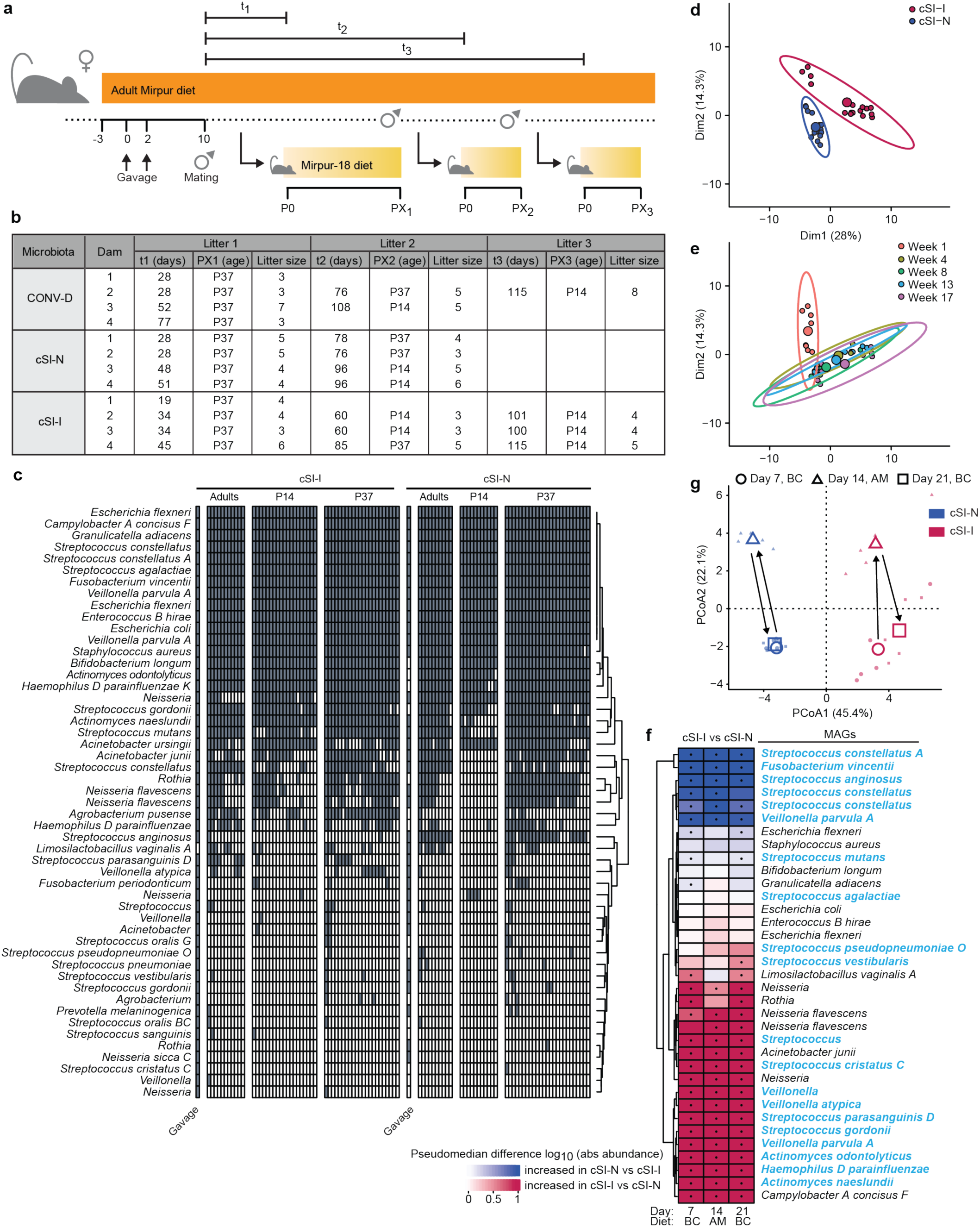
Intergenerational transmission of duodenal bacteria. **a,** Design of the intergenerational transmission experiment. **b,** Summary of animals collected during intergenerational experiment. Litter sizes and the age of offspring at the time of sample collection are shown for each dam (n=4 dams/group). t_1_-t_3_ denotes the number of days between first mating and birth of the litter. **c,** MAGs present at >0.001% relative abundance in gavage mixes, dams, and their P14 and P37 offspring. **d,e,** Principal component analysis of MAGs in feces of dams ordinated by their log_10_ absolute abundance and colored by bacterial consortium (**d**) or timepoint (**e**). The composition of community membership differs significantly between cSI-I and cSI-N dams (**i**, *P*=0.01, PERMANOVA) but is stable after the first week of colonization (**j**, *P*=0.09, PERMANOVA weeks 4 through 17). Each small point represents a fecal sample from one animal. Larger filled circles are the centroids for the ellipses shown, which denote the 95% confidence interval. **f,** Detectable MAGs (log_10_ count > 0) in mice from the ‘Diet oscillation experiment’ (**Extended Data Fig. 1g**) are included in the heatmap; MAGs that match ‘core taxa’ described in children with EED are highlighted in blue, as in Fig. 1b, MAGs whose absolute abundance differs significantly between cSI-I and cSI-N mice are indicated (•, *P*-adj < 0.05, FDR-corrected Wilcoxon rank-sum). **g,** Principal component analysis of MAGs in feces of mice during the transition from breeder chow (the larger shape denotes the centroid of points for each diet and timepoint, circle) to Adult Mirpur (triangle) and back to breeder chow (square).

**Extended Data Fig. 3.**
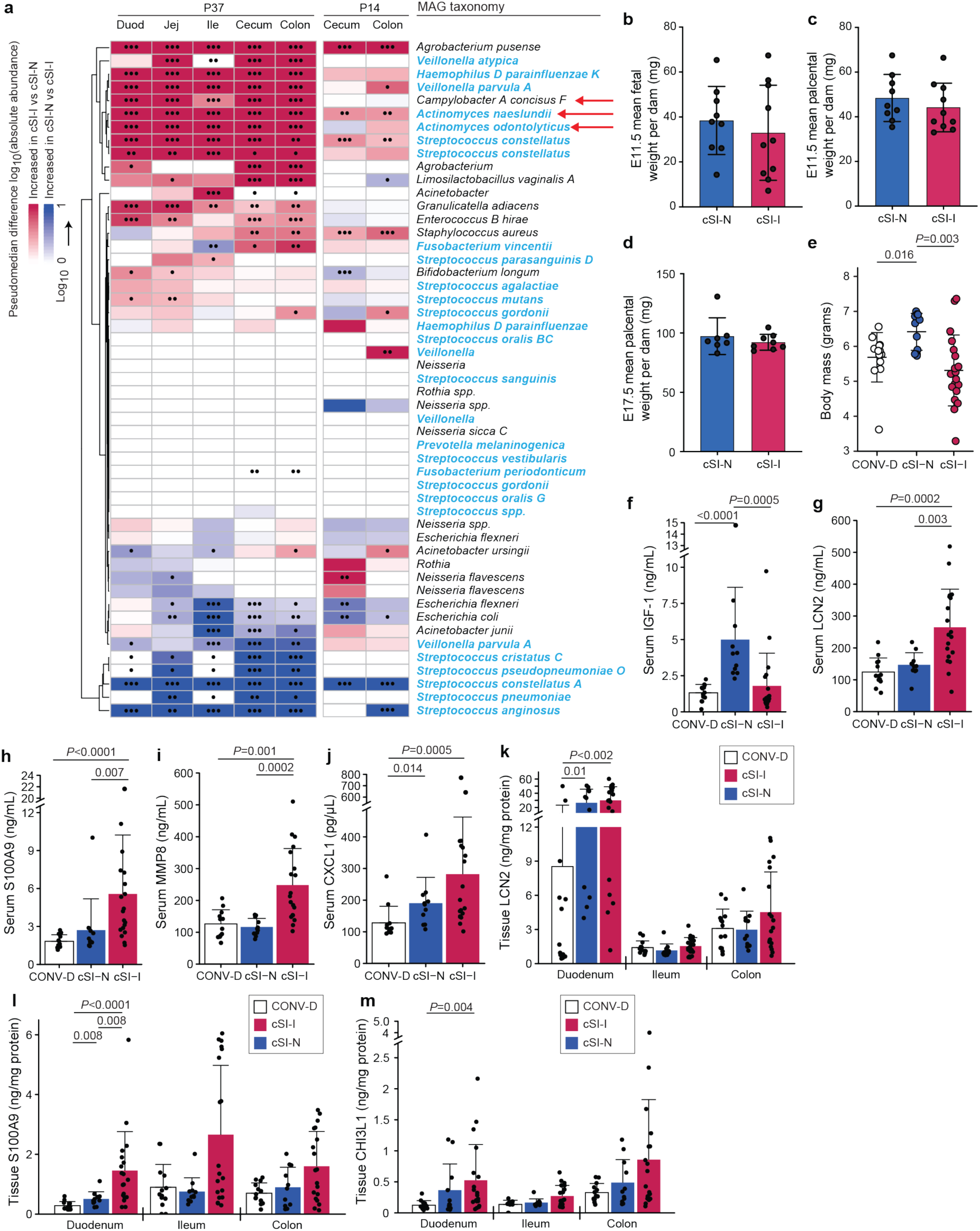
Growth and inflammatory phenotypes of P14 offspring and fetuses. **a,** MAGs whose absolute abundance differs significantly between P14 cSI-I and cSI-N mice are indicated (•, *P*-adj < 0.05, ••, *P*-adj < 0.01, •••, *P*-adj < 0.001, FDR-corrected Wilcoxon rank-sum). All 51 MAGs are included in the heatmap. **b,** Average masses of fetuses per dam at E11.5, n=9-10 dams/group; n=4-8 fetuses/litter. **c,d,** Average masses of placentas within each dam at E11.5 (**c**) and E17.5 (**d**); n=7-10 dams/group; n=2-10 conceptuses/litter. For **b-d**, *P*-values determined with unpaired t-tests. **e,** Body mass of P14 offspring of dams colonized with the cSI-I consortium (red), cSI-N consortium (blue), or cecal contents from a conventional mouse (CONV-D). Each point represents an individual animal. **f,** Serum levels of IGF-1 in P14 offspring. **g-j,** Serum levels of the inflammation-associated proteins, LCN2 (**g**), S100A9 (**h**), MMP8 (**i**), and CXCL1 (**j**), in P14 offspring. **k-m,** Levels of the inflammation-associated proteins LCN2 (**k**), S100A9 (**l**), and CHI3L1 (**m**) in duodenal, ileal, and colonic tissue of P14 animals. For **a,e-m**, n=11-19 mice/group, 3-8 pups/litter, *P*-values were determined using Wilcoxon rank-sum with Tukey’s post-hoc tests. For **b-m**, all panels, mean values ± s.d. are shown.

**Extended Data Fig. 4.**
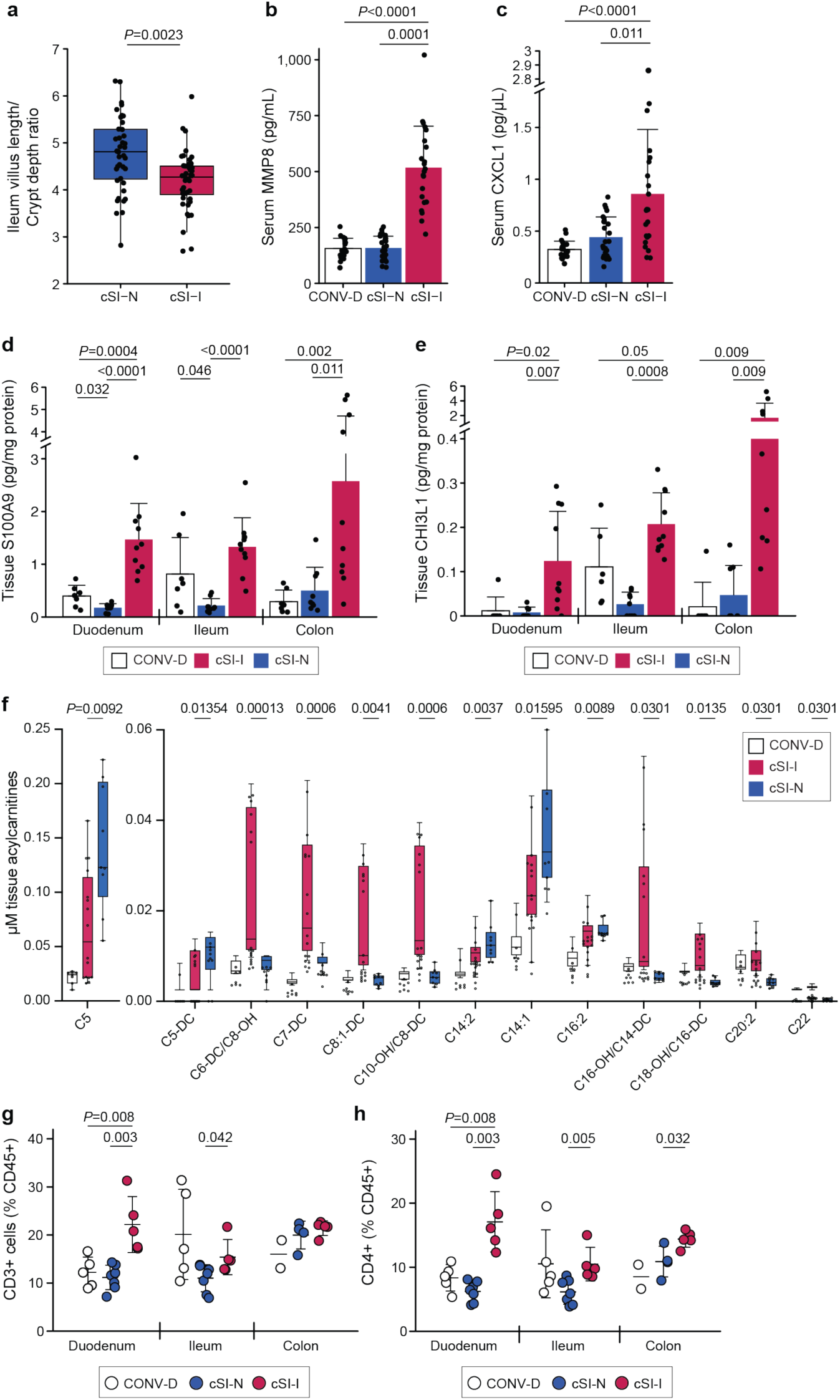
Assessment of EED biomarkers and immune profiling of P37 animals. **a,** Ratio of villus length to crypt depth in the ileums of cSI-I and cSI-N mice (n=5 mice/group, Wilcoxon rank-sum test; boxes denote interquartile range). **b,c,** Levels of serum MMP8 (**b**), and CXCL1 (**c**), in the P37 offspring of C57Bl/6J dams colonized with the cSI-I or cSI-N consortia, or with mouse microbiota harvested from the cecal contents of conventionally raised (CONV-D) C57Bl/6J mice (n=21-25 mice/group). **d,e,** Levels of S100A9 (**d**), and CHI3L1 (**e**), in the duodenal, ileal, and colonic tissue of P37 animals (n=7-10 animals/group). **f,** Concentration of acylcarnitines in small intestinal tissue of P37 animals. Acylcarnitines shown exhibited a statistically significant difference in levels between cSI-I versus cSI-N groups (n=7-16 mice/group, Mann-Whitney tests with Benjamini, Krieger, Yekutieli two-stage step up FDR correction, *q*-values shown). **g,h,** Frequency of CD3^+^ (**g**), and CD4^+^ (**h**) immune cells in duodenal, ileal and colonic lamina propria (n=4-7 mice/group, except for colonic tissue collected from CONV-D animals for which n=2 and a statistical comparison was excluded). For panels **b-e** and **g-h**, bars denote mean ± s.d. and *P*-values were determined by Wilcoxon rank-sum tests.

**Extended Data Fig. 5.**
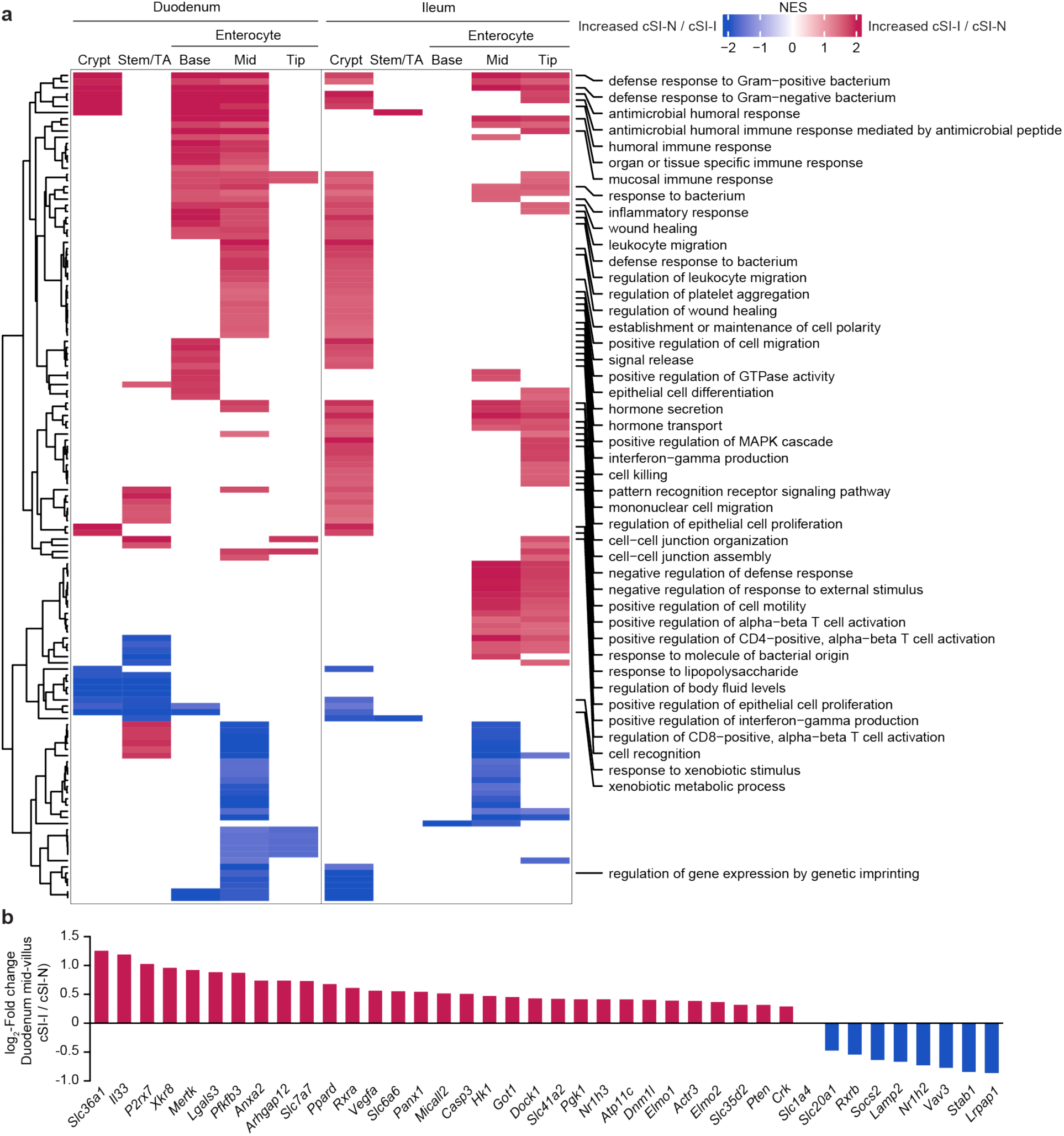
snRNA-Seq differential gene expression in the duodenum and ileum of P37 cSI-I versus cSI-N animals. **a,** Normalized enrichment scores (NES) for all GO categories (rows) that were significantly enriched (*q* < 0.05, GSEA) in at least two of the indicated cell populations (columns) in either the duodenum (left) or ileum (right). Immune-related GO terms are labeled. If a GO category was non-significant, it was assigned an NES value of 0. **b,** Significantly differentially expressed genes comprising the ‘leading edge’ of the efferocytosis gene set in duodenal mid-villus enterocytes (cSI-I versus cSI-N; *P*-adj < 0.05, GSEA). See **Supplementary Table 3c** for annotations.

**Extended Data Fig. 6.**
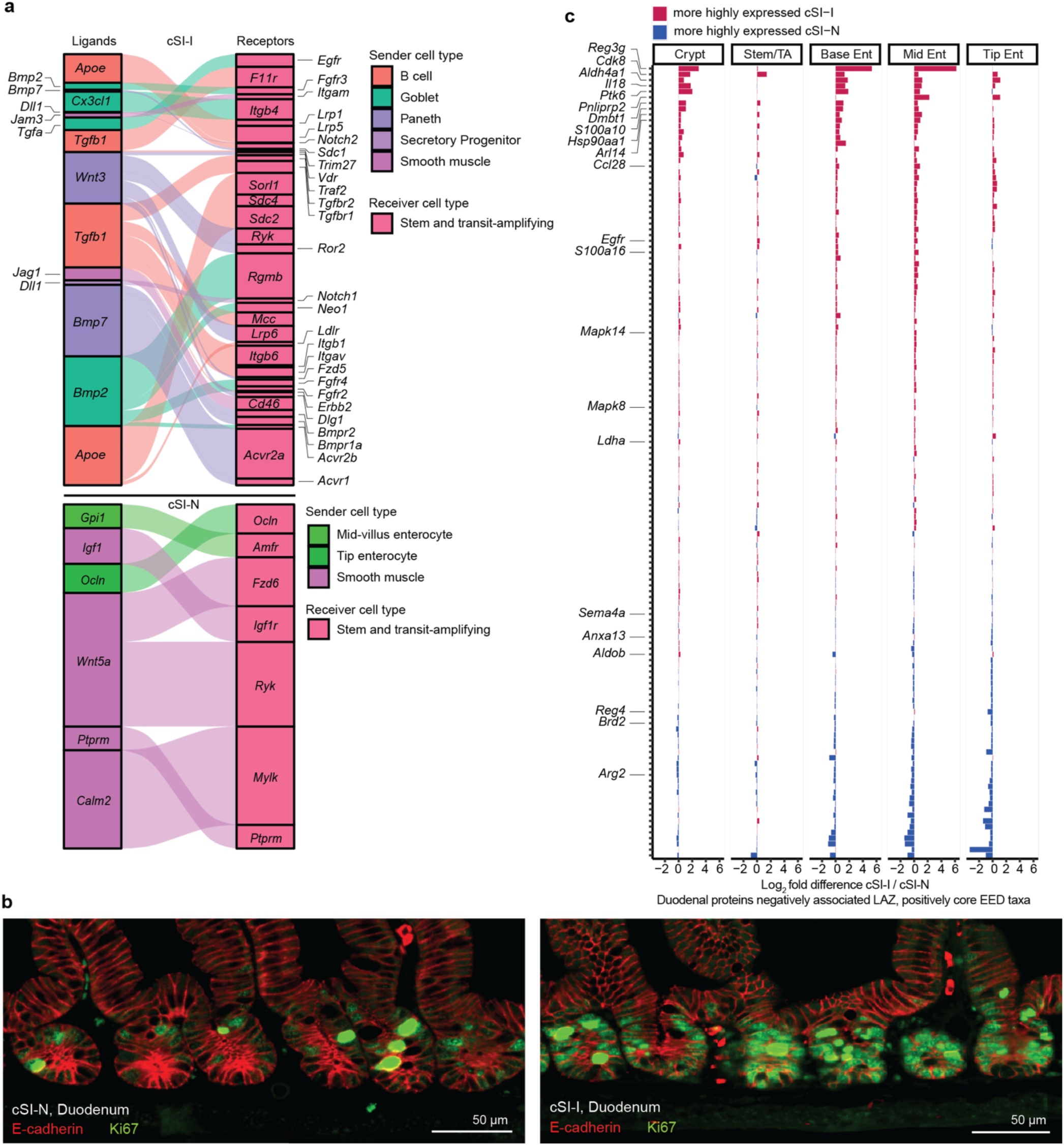
Expression of genes encoding orthologs of proteins quantified in the duodenal mucosa of children with EED and proliferative inter-cellular signaling in P37 animals. **a,** *NicheNet* analysis of the expression of receptors (right) in stem and transit-amplifying (TA) cells for ligands (left) expressed by various cell types in the ileum of P37 animals. Upper panel – ligands expressed more highly in cSI-I animals; lower panel – ligands expressed more highly in cSI-N animals. Boxes are sized proportionally to the weighted ligand/receptor score, related to Fig. 2d. **b,** Representative sections of crypts from the duodenum of cSI-N (left) and cSI-I (right) P37 mice. Sections were stained with antibodies to E-cadherin (red) and Ki67 (green). Scale bar, 50 µm. **c,** Expression of transcripts (rows) in cell populations (columns) in the duodenum of P37 animals that match proteins, quantified in biopsies of the duodenal mucosa of Bangladeshi children with EED, that were (i) negatively correlated with their LAZ and (ii) positively correlated with the absolute abundances of ‘core’ EED bacterial taxa; the abundances of these taxa were positively correlated with their degree of stunting); related to Fig. 2c (see **Supplementary Table 3c** for a list of the mouse homologs of LAZ-associated human proteins).

**Extended Data Fig. 7.**
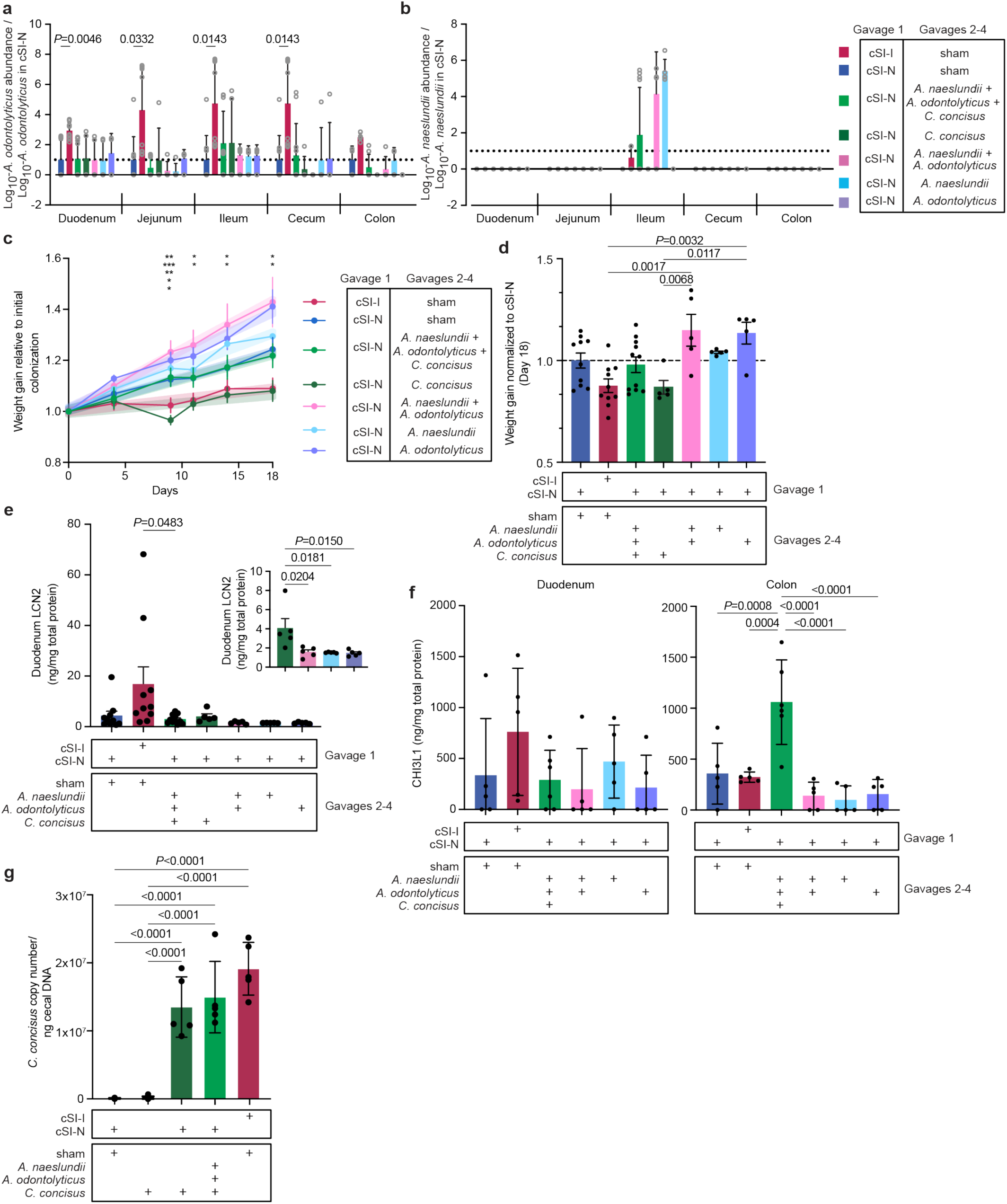
Abundance of *Actinomyces* in isolate ‘add-in’ experiments and. **a,b,** Mice were initially colonized with either the cSI-N or cSI-I consortium (‘gavage 1’). Gavages 2-4 were performed on experimental days 9 through 11, as in Fig. 4a. The absolute abundances of *A. odontolyticus* isolate Bg044 (corresponding to MAG044, panel **a**) and *A. naeslundii* isolate Bg041 (corresponding to MAG041, panel **b**), shown relative to their absolute abundance in the cSI-N/sham-gavaged controls (a ratio of 1 is indicated by dotted line; mixed-effects analysis with Dunnett’s multiple comparisons to cSI-N/sham control; mean values ± s.d. are shown.) **c,** Changes in body mass relative to time of colonization in mice in ‘add-in’ experiments (points and error bars denote mean values ± s.e.m.; shaded, thicker line represents results of linear regression. \**P*<0.05, \*\**P*<0.01, \*\*\**P*<0.001 for Tukey’s multiple comparisons demonstrating significant differences between *C. concisus*-gavaged animals compared to cSI-N/mock, cSI-N/*A.naeslundii*, *A. odontolyticus*, or *Actinomyces* from two-way repeated measures ANOVA; all stats are reported in **Supplementary Table 2b**). **d**, Relative weight gain in isolate-gavaged and cSI-I/mock-gavaged animals relative to weight gain in cSI-N mice (one-way ANOVA with Tukey’s post-hoc, mean ± s.e.m.). **e,** Levels of LCN2 protein in duodenal tissue on experimental day 18 (one-way ANOVA with Tukey’s multiple comparisons; mean values ± s.e.m. are shown). The inset shows comparisons between individual isolates. **f,** Tissue levels of CHI3L1 (Chitinase-3-like protein 1) in the duodenum and colon (one-way ANOVA with Tukey’s multiple comparisons, n=5-6 mice/group, mean ± s.d. shown). For **a-e**, n=5-11 mice/group combined across two independent experiments. **g,** Abundance of *C. concisus* in cecal contents of mice mono-colonized (gavaged five times on days 0, 3, 5, 7, and 14) compared to ‘add-in’ experiment (n=5-7 mice/group, mean ± s.d. shown, one-way ANOVA with Tukey’s multiple comparisons).

**Extended Data Fig. 8.**
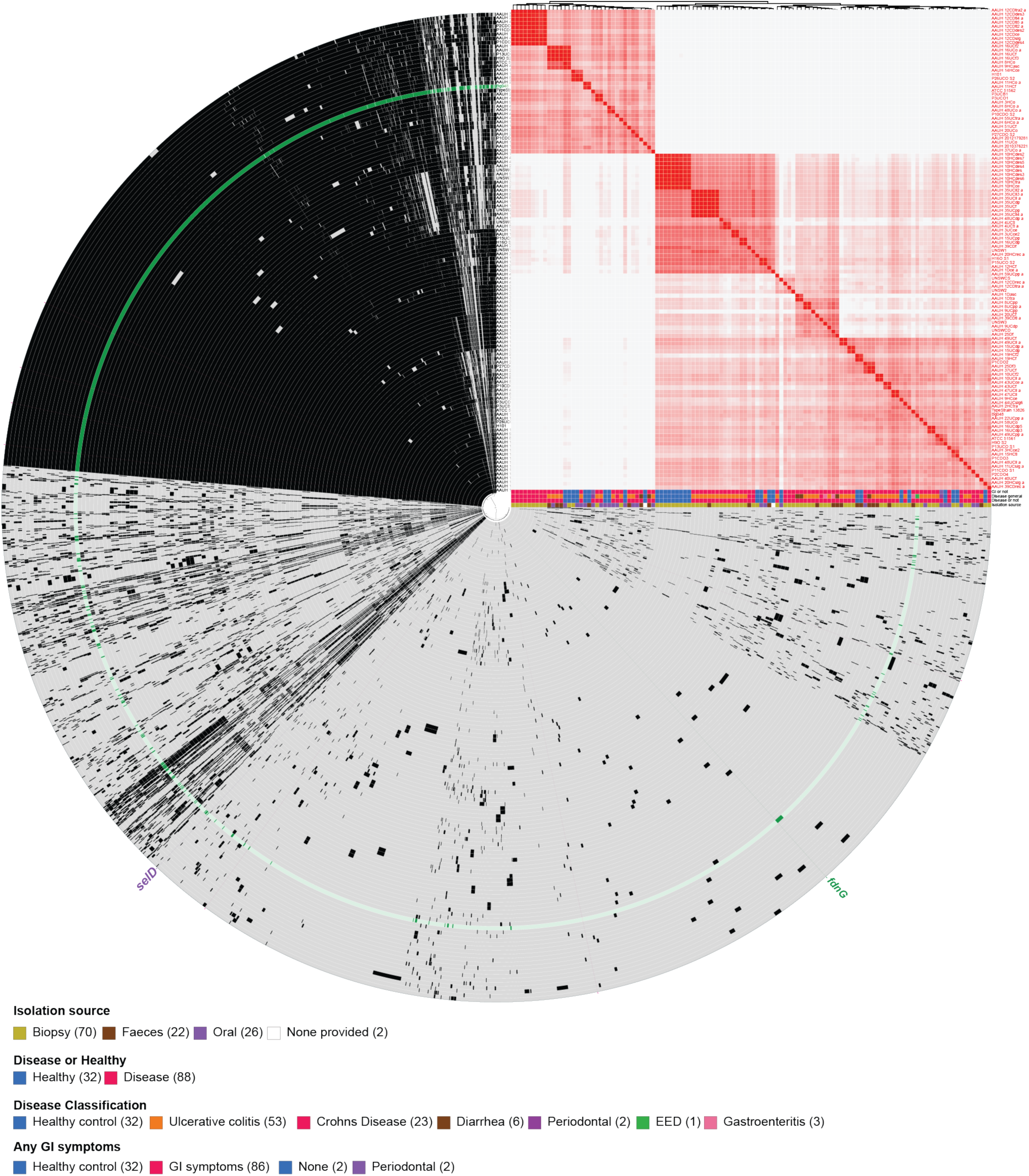
Comparative genomic analysis of *C. concisus* isolates. 120 whole *C. concisus* genomes are included in the circle phylogram that were isolated from intestinal biopsies, feces, or saliva of healthy or diseased individuals. Genes that are present are shown in black; if genes are absent, in gray. The Bangladeshi isolate from children with EED (Bg048) is highlighted in green. The nitrate-inducible formate dehydrogenase (*fdnG*) unique to this Bangladeshi isolate and a selenate di-kinase (*selD*) unique to the Bangladeshi isolate and one other genome are highlighted. Heatmap shows ANI comparisons between all isolates.

**Extended Data Fig. 9.**
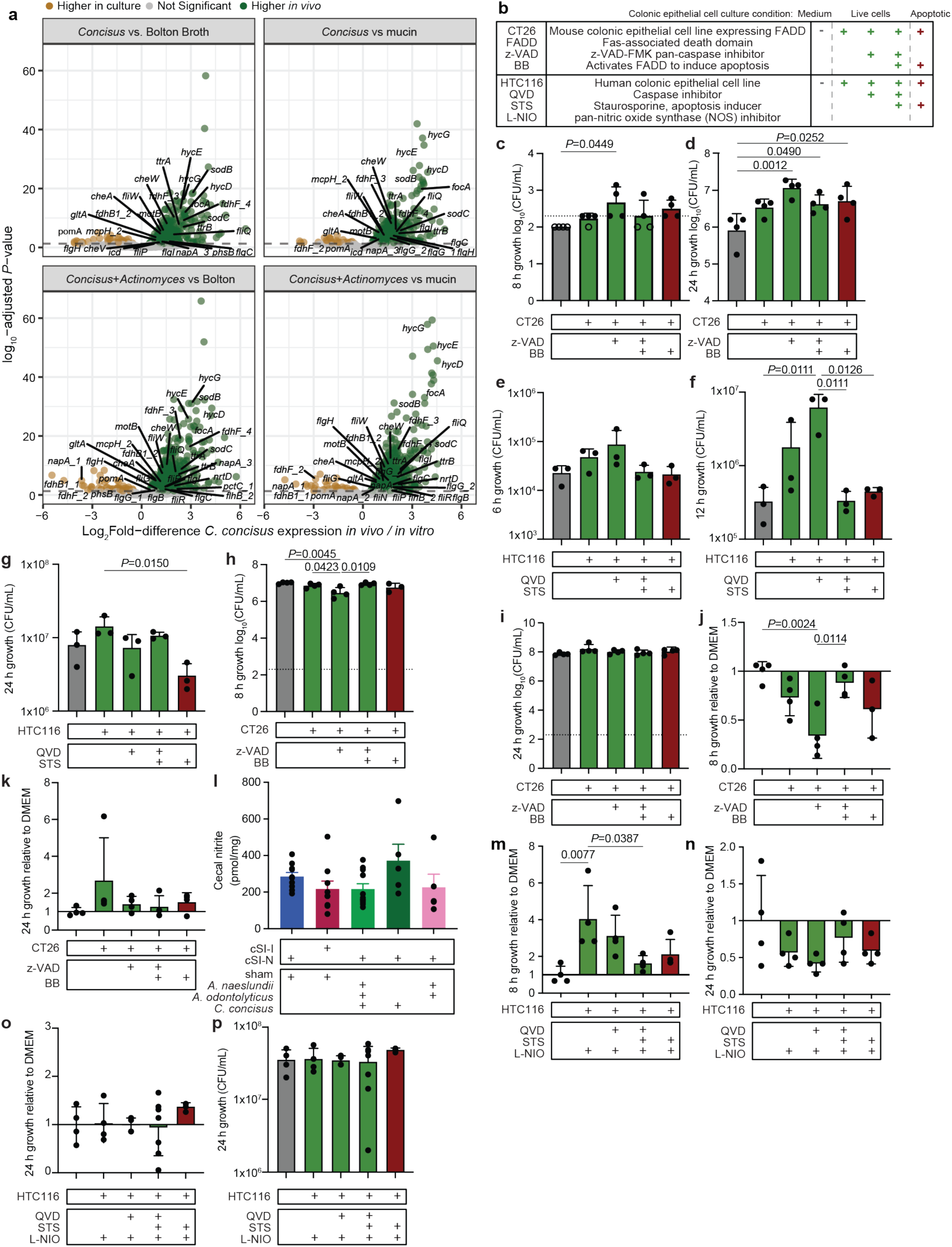
*C. concisus* growth in spent medium collected from colonic epithelial cells. **a,** *C. concisus* gene expression in the gut (*Concisus*, *Concisus*+*Actinomyces*) from the isolate ‘add-in’ experiment (Fig. 5) compared to growth in rich medium (Bolton broth, or Bolton broth supplemented with 1% mucin). Green, *P*-adj < 0.05 and higher expression in the gut; brown, *P*-adj < 0.05 and higher expression *in vitro* (*DESeq2* Wald test, n=4 biological replicates per condition). Genes involved in formate metabolism and anaerobic respiration are labelled. **b,** Abbreviations of cell lines and treatments used to generate conditioned medium from mouse and human colonic epithelial cells that were live or apoptotic. *C. concisus* growth in conditioned medium acquired from live cells are colored in green; apoptotic cells, in red. QVD, Quinoline-Val-Asp-Difluorophenoxymethylketone (caspase inhibitor); L-NIO, *N*^5^-(1-Iminoethyl)-L-ornithine, dihydrochloride (nitric oxide synthase inhibitor). **c,d,** Microaerophilic growth of *C. concisus* after 8 (**c**) or 24 hours (**d**) in conditioned medium from mouse CT26 colonic epithelial cells that were live (CT26:FADD, z-VAD, z-VAD+BB) or treated with an inducer of apoptosis (BB). Points shown with an open rather than filled circle denote abundances that were below the limit of detection at the time of quantification and are presented at the limit of detection of the assay. **e-g,** Microaerophilic growth of *C. concisus* after 6 (**e**), 12 (**f**), or 24 (**g**) hours in spent medium from HTC116 human colonic epithelial cells that were live (QVD, QVD+STS) or that had been treated with an inducer of apoptosis (STS). **h-k,** Anaerobic growth of *C. concisus* after 8 (**h, j**) or 24 hours (**i, k**) in conditioned medium from live CT26 mouse colonic epithelial cells (CT26:FADD, z-VAD, z-VAD+BB) or from cells treated with an inducer of apoptosis (BB). Panels **j** and **k** show growth relative to (as a ratio to) growth in tissue culture medium. **l,** Levels of nitrite in cecal contents 9 days post-isolate gavage (n=5-11 mice/condition, combined across two independent experiments). **m,n,** Microaerophilic growth of *C. concisus* after 8 (**m**) or 24 (**n**) hours growth in conditioned medium from HTC116 human colonic epithelial cells acquired from live cells (QVD, QVD+STS) or cells treated with an inducer of apoptosis (STS). All cells were treated with an inhibitor of nitric oxide synthase (L-NIO). Growth shown as a ratio to growth in cell culture medium alone. **o,p,** Anaerobic growth of *C. concisus* after 24 hours of incubation in conditioned medium from HTC116 cells collected from live cells (QVD, QVD+STS) or cells that had been treated with an inducer of apoptosis (STS). All cells were treated with an inhibitor of nitric oxide synthase (L-NIO). Panel **o** shows growth as a ratio to growth in tissue culture medium alone; **p**, total bacterial growth. For **c-p**, bars denote mean ± s.d., *P-*values determined with one-way ANOVA and Tukey’s multiple comparisons. For **c,d**, **g-k**, **m-p**, n=4 biological replicates/condition; for **e,f**, n=3 biological replicates/condition.

**Extended Data Fig. 10.**
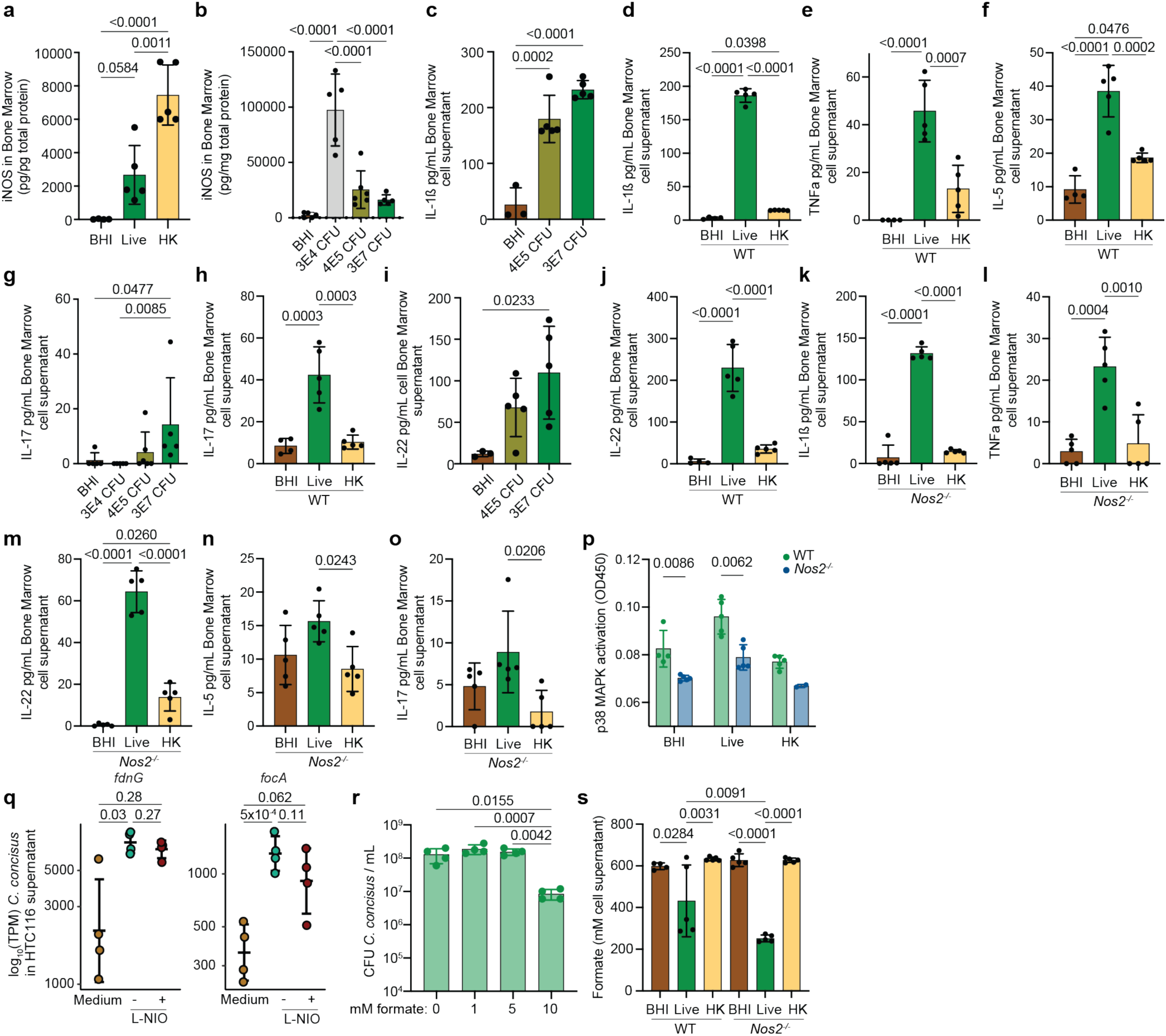
*C. concisus* induces iNOS and immune signaling. **a,b,** Bone marrow cells from wild-type (WT) mice were incubated for 48-hours with fresh (Live) or heat-killed (HK) *C. concisus* culture, or bacterial medium as a negative control. **b,** Bone marrow cells were co-cultured for 48-hours with increasing amounts of live *C. concisus* culture. Proteins were extracted from bone marrow cell pellets and iNOS levels were quantified. **c-j,** Bone marrow cells collected from wild-type (WT) mice were incubated for 48 hours with fresh *C. concisus* culture (Live, CFU), heat-killed *C. concisus* culture (HK), or bacterial medium as a negative control (BHI). IL-1ß (**c,d**), TNFa (**e**), IL-5 (**f**), IL-17 (**g,h**), and IL-22 (**i,j**) were quantified in cell supernatant. **k-o**, Bone marrow cells collected from *Nos2^−/−^* mice were incubated for 48 hours with fresh (Live) or heat-killed (HK) *C. concisus* culture. Again, IL-1ß (**k**), TNFa (**l**), IL-22 (**m**), IL-5 (**n**) and IL-17 (**o**) were quantified from cell culture supernatants. **p,** p38 MAPK activation was assessed in bone marrow cell lysates after 48 hours incubation bacterial medium (BHI), fresh *C. concisus* culture (Live), or heat-killed (HK) *C. concisus* culture (multiple unpaired t-tests with two-stage step-up FDR correction). **q,** Expression of the *C. concisus* nitrate-inducible formate dehydrogenase (*fdnG*) and formate transporter (*focA*) when cultured in supernatants collected from HTC116 cells that were or were not treated with the NOS inhibitor L-NIO or cell culture medium alone (statistics shown are unadjusted *P* values from the *DESeq2* Wald test, transcripts per million). **r,** *C. concisus* was cultured in Bolton broth supplemented with various levels of formate; growth was quantified after 24 hours. **s,** Formate was quantified in supernatants from bone marrow cells incubated for 48 hours with bacterial culture medium (BHI), fresh *C. concisus* culture (Live), or heat-killed *C. concisus* culture (HK). For all panels, n=4-5 biological replicates/condition, bars denote mean ± s.d.. For **a-o**, **r**,**s**, *P*-values shown are the result of one-way ANOVA followed with Tukey’s multiple comparisons.

## SUPPLEMENTARY TABLES

**Supplementary Table 1. Diet and bacterial abundance information from gnotobiotic mouse experiments. a,** Diets used in gnotobiotic mouse studies. Ingredients in and nutritional analysis of each diet. **b,** Cultured isolates and taxonomic classification from BEED study duodenal aspirates. **c,** MAG assembly information and MAG taxonomic classification. (i), Assembly and taxonomic classification. (ii), Sample metadata. **d,** Abundances of MAGs from gnotobiotic animal experiments. (i), Diet oscillation experiment, log_10_-tpm (**Extended Data Figs. 2f,g**). (ii), Intergenerational transmission experiment, absolute abundance: P37 animals (**Figs. 1c, 3j, Extended Data Figs. 2c, 3a**), P14 animals (**Extended Data Fig. 3a**) and adult mice (**Extended Data Figs. 2c-e**). (iii), Cohousing experiment, absolute abundance (**Figs. 3h,i**). (iv), Isolate add-in experiments, absolute abundance (**Fig. 4b, Extended Data Figs. 7a,b**). Abundance values are log_10_-transformed. *P*-adj values determined by Wilcoxon rank-sum tests and FDR correction. **e,** *C. concisus* abundance in (i) mono-colonization and (ii) add-in experiments assessed by qPCR.

**Supplementary Table 2. Phenotypic data from gnotobiotic mouse experiments. a,** Body weights from gnotobiotic animal experiments. (i) Preliminary test of consortia in recently weaned mice (**Extended Data Fig. 1a**); (ii), Intergenerational transmission experiment: P37 animals, P14 animals (**Extended Data Fig. 3e**) and conceptuses (**Fig. 1c, Extended Data Figs. 3b-d**) (Wilcoxon rank-sum tests). (iii), Cohousing experiments (linear mixed effects modeling). (iv), Gavage of pathology-associated isolates, ‘add-in’ experiments (**Extended Data Figs. 7c,d**) (repeated measures ANOVA). **b,** Protein levels in tissues from gnotobiotic animal experiments. (i), First test of consortia in recently weaned mice (**Extended Data Figs. 1b,c,f**). (ii), Diet oscillation experiment (**Extended Data Fig. 1h**). (iii), Intergenerational transmission experiment: P37 duodenum, ileum, colon, and serum (**Figs. 1f-h, Extended Data Figs. 4b-e**); P14 duodenum, ileum, colon and serum (**Extended Data Figs. 3f-m**); adult duodenum, ileum, colon, and serum. (iv), Cohousing experiments: duodenum, ileum, colon and serum (**Figs. 3b-c**). (v), Isolate add-in experiments: duodenum, colon and serum (**Fig. 4c-e, Extended Data Figs. 7e-f**). (vi), *C. concisus* mono-colonization experiment: duodenum, ileum, colon and serum. **c,** Histomorphometric analysis of small intestines of P37 animals from the intergenerational transmission experiment. (i), Duodenal (**Figs. 1d,e**) and (ii) ileal villus height, crypt depth, villus to crypt ratio (**Extended Data Fig. 4a**), and Ki67 quantification (**Fig. 2d, Extended Data Fig. 6b**). *P*-values determined using Tukey’s post-hoc tests. **d,** Micro-computed tomography of femurs of P37 animals from the intergenerational experiment. *P*-values were determined using Wilcoxon rank-sum tests. **e,** Quantification of small intestinal acylcarnitines in P37 offspring (log_10_-µM) (**Extended Data Fig. 4f**). **f,** Immune cell profiling of tissues from gnotobiotic animal experiments. (i), Intergenerational transmission experiment: P37 duodenum, ileum, colon, spleen and meninges (**Figs. 1i,j, Extended Data Figs. 4g,h, Supplementary Figs. 1a-c**); duodenum, ileum, spleen and meninges of the dams. (ii), Cohousing experiment: bulk small intestinal tissue, spleen and meninges (**Figs. 4d-g**). *P*-values determined using Wilcoxon rank-sum tests.

**Supplementary Table 3. Supplemental data for host RNA-sequencing. a,** Bulk tissue RNA-Seq from gnotobiotic animal experiments. (i), Preliminary tests of consortia in recently weaned animals, significantly differentially expressed genes and results of GSEA between cSI-I and cSI-N-colonized mice 28 days after gavage in the duodenum, jejunum and ileum (**Extended Data Figs. 1d,e**). (ii), Intergenerational transmission experiment: gene set enrichment analysis (GSEA) of gene expression in the duodenum, ileum and colon of P37 animals. (iii), Cohousing experiment: differentially expressed genes in duodenal samples for: non-cohoused cSI-I controls versus non-cohoused cSI-N controls and cohoused versus non-cohoused cSI-N controls. There were no statistically significant differentially expressed genes between cohoused and cSI-I controls. GSEA of gene expression in the colon of non-cohoused cSI-I controls vs. non-cohoused cSI-N controls and cohoused vs. non-cohoused cSI-N controls (Fig. 4f). (iv), Add-in experiment: GSEA of gene expression in the colon. Pathways shown were enriched in all the following conditions: cohoused animals vs. cSI-N non-cohoused controls, *C. concisus*+*Actinomyces*-gavaged animals vs. cSI-N sham-gavaged controls and *C. concisus*-gavaged animals vs. cSI-N sham-gavaged controls. Values provided (*P-* and *q-*values, enrichment scores, rank, leading edge, core enrichment) are for *C. concisus+Actinomyces*-gavaged vs. cSI-N sham-gavaged animals (Fig. 4f). **b,** Proportions of cell types quantified by snRNA-Seq in the duodenums (i) and ileums (ii) of P37 animals from the intergenerational transmission experiment (**Figs. 2a,b**) (n=3 mice/group, statistical significance determined by *edgeR*). **c,** snRNA-Seq analysis of duodenums and ileums of P37 offspring from intergenerational transmission experiment. (i) Pseudo-bulk differential expression of enterocyte clusters in the duodenum and ileum of cSI-I compared to cSI-N animals (n=3 mice/group; *DESeq2* Wald test). (ii) GSEA of gene expression along the crypt-villus axis (**Extended Data Fig. 5a**). (iii) GSEA and annotations for genes involved in efferocytosis (**Extended Data Fig. 5b**). (iv) ‘Differential *NicheNet’* results (**Fig. 2c, Extended Data Fig. 6a**). (v) GSEA of homologues in the mouse gut (P37 duodenum and ileum) to LAZ-associated proteins quantified in the duodenal mucosa of children in the BEED study (**Fig. 2e, Extended Data Fig. 6c**). **d,** GSEA results in the duodenum and colon for bulk RNA-seq datasets of animals from cohousing and ‘add-in’ experiments; all treatment groups presented were compared to their cSI-N counterparts (Fig. 4f).

**Supplementary Table 4. Comparative genomics analysis. a,** Isolate genome assembly information and taxonomic classification. (i), Quality assessment and taxonomic classification of assembled isolate genomes. (ii), Average nucleotide identity (ANI) of isolate genomes with their related MAGs. (iii), Species-specific virulence factors in MAGs, their respective isolate genomes, and related reference genomes. **b,** *In silico* metabolic reconstructions of MAGs and isolate genomes. (i), mcSEED-based metabolic annotation for each MAG by gene. (ii), Annotations for pathology-associated MAGs and phylogenetically related organisms by gene. (iii) ‘Metabolic phenotype’ prediction key. **c,** *C. concisus* pan-genome analysis (**Extended Data Fig. 8**). (i), List of 120 isolate genomes included in pan-genomic analysis and their association with disease. Enrichment analyses performed on predicted genes (ii), COG20 functional annotations (iii) and COG Pathway annotations (iv). Enrichment results were included if functional groups were enriched in isolates from diseased conditions (Crohn’s Disease, Ulcerative colitis, Gastroenteritis, Periodontitis) including EED, but not in isolates from healthy controls.

**Supplementary Table 5. Supplemental data for microbial RNA-sequencing. a,** Significantly differentially expressed *C. concisus* genes in the gut compared to growth in culture (**Extended Data Fig. 9a**) (*DESeq2* Wald test). (i,ii), *C. concisus* gene expression in the gut (single isolate ‘add-in’) compared to rich medium (Bolton broth ± 1% mucin). (iii,iv) *C. concisus* gene expression when gavaged concurrently with the *Actinomyces* strains (*C. concisus*+*Actinomyces*) compared to gene expression in rich medium (Bolton broth ± 1% mucin). **b,** *C. concisus* gene expression in the mouse gut in conditions where inflammation was produced [*C. concisus* (i), *C. concisus*+*Actinomyces* (ii), cSI-I (iii)] compared to cSI-N/mock-gavaged controls (**Figs. 5b,c**). **c,** *C. concisus* gene expression in spent medium collected from HTC116 cells compared to medium alone (Fig. 5l) (i). *C. concisus* gene expression in conditioned medium from HTC116 cells treated with the pan-nitric oxide inhibitor L-NIO compared to medium control (ii).

**Supplementary Table 6. Antibody panels for flow cytometry analyses.** Antibodies used for flow cytometry. (i), Antibodies used for flow cytometry of intestinal immune cells in intergenerational transmission and cohousing experiments (**Figs. 1i,j, Extended Data Figs. 4g,h**). (ii and iii), Antibodies used for flow cytometry of splenic and meningeal immune cells in intergenerational transmission (ii) and cohousing experiments (iii, **Figs. 3f,g**).

**Supplementary Table 7. Metadata associated with sequencing datasets.** Sequencing metadata from gnotobiotic animal and bacterial culture experiments. **a,** Bulk RNA-Seq collected from preliminary test of consortia (**Extended Data Figs. 1d,e**). **b,** Intergenerational transmission experiment: (i) short read shotgun DNA sequencing (**Figs. 1b,3j, Extended Data Figs. 2c-e, 3a**), (ii) snRNA-Seq (**Figs. 2a-c,e, Extended Data Figs. 5, 6a,b**), (iii) bulk tissue RNA-Seq (Fig. 4f) and (iv) microbial RNA-Seq. **c,** Cohousing experiment: (i) short read shotgun DNA sequencing (**Figs. 3h,i**), (ii) bulk tissue RNA-Seq (Fig. 4f) and (iii) and microbial RNA-Seq. **d,** Isolate add-in experiments: (i) short read shotgun DNA sequencing (**Fig. 4b, Extended Data Figs. 7a,b**), (ii) and bulk tissue RNA-Seq and (iii) microbial RNA-Seq. **e,** *C. concisus* microbial RNA-Seq in culture medium and conditioned medium collected from HTC116 cells (**Fig. 5l, Extended Data Fig. 9a**).

**Supplementary Fig. 1.**
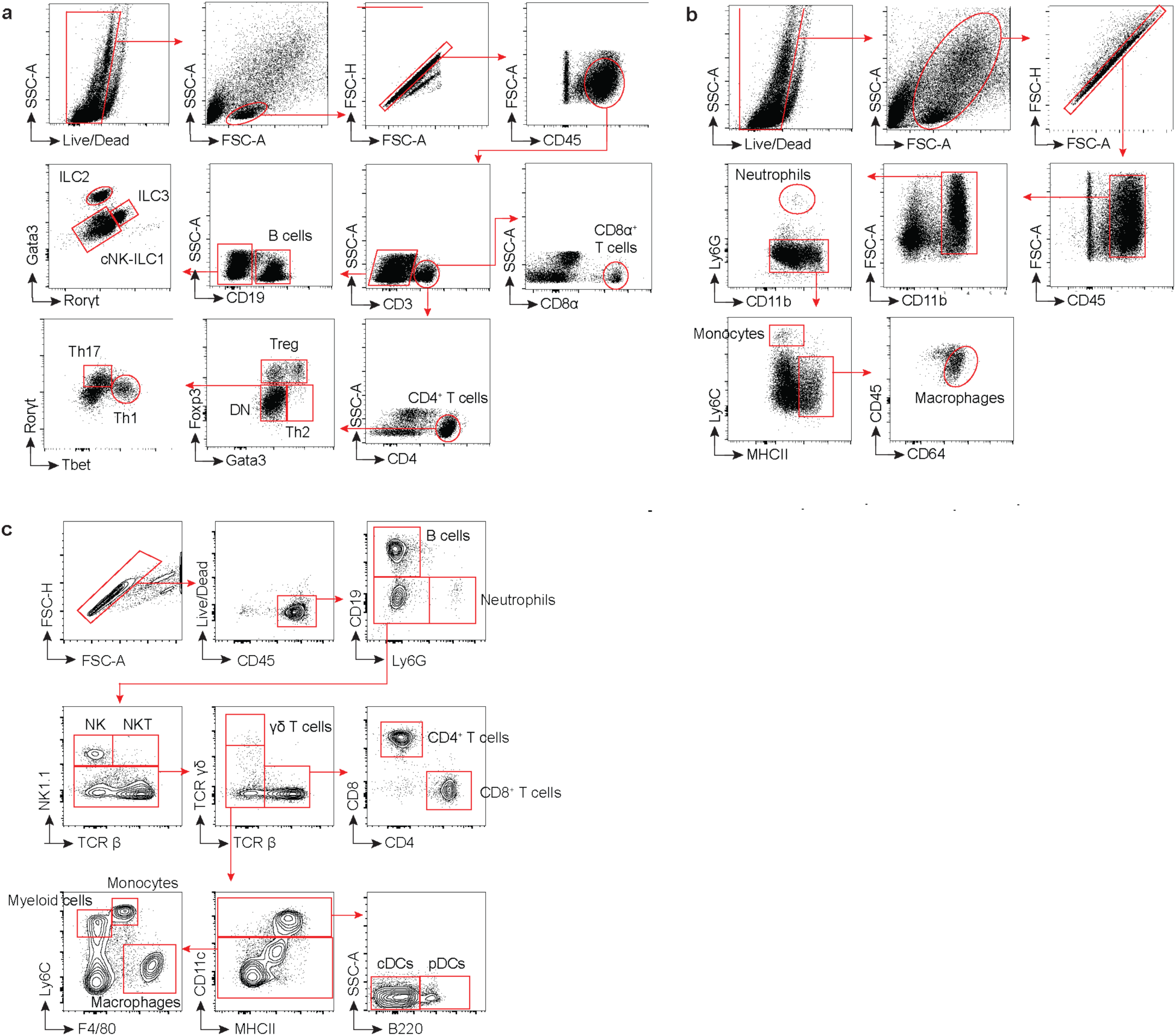
Gating strategies for flow cytometry. **a,b,** Representative gating strategy for quantifying lymphoid (**a**) and myeloid (**b**) immune cells in the intestinal lamina propria. **c,** Representative gating strategies for meninges and spleen. See **Supplementary Table 6** for antibodies used.

